# Nub1 traps unfolded FAT10 for ubiquitin-independent degradation by the 26S proteasome

**DOI:** 10.1101/2024.06.12.598715

**Authors:** Connor Arkinson, Ken C. Dong, Christine L. Gee, Shawn M. Costello, Susan Marqusee, Andreas Martin

## Abstract

The ubiquitin-like modifier FAT10 targets hundreds of proteins in the mammalian immune system to the 26S proteasome for degradation. This degradation pathway requires the cofactor Nub1, yet the underlying mechanisms remain unknown. Here, we reconstituted a minimal *in vitro* system and revealed that Nub1 utilizes FAT10’s intrinsic instability to trap its N-terminal ubiquitin-like domain in an unfolded state and deliver it to the 26S proteasome for engagement, allowing the degradation of FAT10-ylated substrates in a ubiquitin– and p97-independent manner. Through hydrogen-deuterium exchange, structural modeling, and site-directed mutagenesis, we identified the formation of a peculiar complex with FAT10 that activates Nub1 for docking to the 26S proteasome, and our cryo-EM studies visualized the highly dynamic Nub1 complex bound to the proteasomal Rpn1 subunit during FAT10 delivery and the early stages of ATP-dependent degradation. These studies thus identified a novel mode of cofactor-mediated, ubiquitin-independent substrate delivery to the 26S proteasome that relies on trapping partially unfolded states for engagement by the proteasomal ATPase motor.

## Introduction

Ubiquitin-mediated substrate degradation by the 26S proteasome relies on a bipartite signal consisting of a suitable ubiquitin modification, like a polyubiquitin chain, and a substrate’s unstructured initiation region that is of sufficient length (at least 20-25 residues) for engagement by the proteasomal AAA+ (ATPases Associated with diverse cellular Activities) motor ^1–4^. Ubiquitinated substrates that lack such an intrinsic unstructured region can be prepared for proteasomal engagement through processing by the AAA+ protein unfoldase p97, also known as Cdc48 in yeast ^5^, which initiates substrate unfolding on a ubiquitin moiety within the ubiquitin chain and releases partially or completely unstructured proteins for degradation by the proteasome ^6–8^. The 26S proteasome is composed of the 20S core particle (CP), which contains a barrel-shaped degradation chamber with sequestered proteolytic active sites, and the 19S regulatory particle (RP) that binds to either end of the CP ^9^. The RP includes three main ubiquitin-receptor subunits, Rpn10, Rpn1, and Rpn13 ^10–13^, the deubiquitinase Rpn11, and the heterohexameric AAA+ motor, which consists of the ATPase subunits Rpt1-Rpt6 ^9,14,15^. After ubiquitin-chain binding to a proteasomal receptor, the ATPase motor engages a substrate’s unstructured initiation region through conserved pore loops in its central channel and uses ATP-hydrolysis-driven conformational changes for mechanical substrate unfolding and translocation into the CP for cleavage ^4^, while Rpn11 catalyzes the co-translocational *en-bloc* removal of ubiquitin modifications ^16–18^.

Recently, several new pathways for proteasomal substrate delivery in ubiquitin-independent manners have been described ^19–21^, yet the underlying principles remain largely elusive. Here, we determine the mechanism for the ubiquitin-independent degradation of substrates carrying the ubiquitin-like (UBL) modifier FAT10 in a novel mode of recruitment to the 26S proteasome. FAT10 (human leukocyte antigen-F-Adjacent Transcript 10) is expressed predominantly in cells of the immune system and controls numerous cellular processes, including apoptosis and antigen presentation ^22–26^. FAT10 can be induced by virus infections or pro-inflammatory cytokines such as TNF-α and INF-γ ^22,24^, and it is prevalent in multiple types of cancers where it aids proliferation and metastasis formation ^27–29^. It regulates hundreds of proteins in their function and abundance by forming non-covalent or covalent interactions ^30^. For covalent attachment, FAT10 is typically conjugated via its C-terminal glycine to lysine residues of substrates that are then targeted for proteasomal degradation ^22,31–33^. Unlike ubiquitin, FAT10 is not removed and recycled, but functions as both the targeting signal and a probable initiation region for degradation ^34^. In its free and conjugated forms, FAT10 is rapidly degraded by the 26S proteasome, with an estimated half-life in cells of ∼1 hour ^31^. It contains two ubiquitin-like domains (UBLs) connected by a short linker, and while there is evidence for its degradation being ubiquitin-independent ^31,35,36^, other studies indicated that turnover predominantly occurs through ubiquitin targeting ^37^. Interestingly, FAT10 does not contain any disordered segments long enough for proteasomal engagement and its degradation *in vivo* is not blocked by p97 inhibitors ^34^, leaving the question of how it can bypass the requirement for a bipartite degradation signal that includes an unstructured initiation region in addition to a targeting signal.

The inflammation-induced protein NEDD8-ultimate-buster-1 (Nub1) and its longer isoform, Nub1L, were shown to bind to and accelerate the degradation of FAT10 ^38,39^. While FAT10 is exclusively found in mammals, Nub1 variants are also present in flies and plants, suggesting a more conserved function that is potentially linked to Nub1’s roles in accelerating NEDD8 degradation ^40^. Nub1 contains an N-terminal UBL domain and three C-terminal ubiquitin-associated domains (UBA1-UBA3), which were proposed to bind to the 26S proteasome and may be responsible for the Nub1-dependent acceleration of FAT10 degradation ^36,41,42^. While Nub1’s UBA domains appear critical for FAT10 binding, they were claimed to be dispensable for facilitating FAT10 degradation, and it was suggested that a ternary complex with the 26S proteasome, yet lacking a direct FAT10-Nub1 interaction, may be sufficient to drive FAT10 turnover ^41^. The proteasomal ubiquitin receptors Rpn10 and Rpn1 were both postulated to bind Nub1’s UBL domain, while Rpn10 was also assumed to bind to Nub1’s UBA domains and FAT10’s UBL2 domain, suggesting some form of competing interactions or order of events that led to a confusing model for FAT10 recruitment ^36,42,43^. It therefore remained completely unclear how FAT10 and Nub1 interact with each other and with the 26S proteasome, and how Nub1 can accelerate the degradation of FAT10.

Here, we *in vitro* reconstituted the ubiquitin-independent degradation of FAT10 by the human 26S proteasome and determined Nub1 as an essential cofactor for both the delivery of FAT10 to the proteasome and its preparation for engagement by the AAA+ motor. Using hydrogen-deuterium exchange with detection by mass spectrometry (HDX-MS), AlphaFold modeling, biochemical assays, and site-directed mutagenesis, we show that Nub1 is an ATP-independent chaperone that ‘traps’ partially unfolded FAT10 in an internal channel between its helical core and UBA domains, and positions FAT10’s N-terminus for insertion into the proteasome. Furthermore, we revealed that FAT10 binding induces an ‘open’ Nub1 conformation with Nub1’s UBL domain undocked from the trap domain. Our cryo-electron microscopy (cryo-EM) studies captured the 26S proteasome during Nub1-dependent FAT10 processing and show a highly flexible Nub1 that specifically interacts through its UBL domain with the T2 site of the proteasome’s Rpn1 receptor subunit. These data thus provide the first mechanistic insight into how a shuttle factor can accelerate the turnover of its target substrates in a ubiquitin-independent manner and, more specifically, explain how Nub1 allows FAT10-modified proteins to bypass p97 requirements for engagement and degradation by the 26S proteasome.

## Results

### Proteasomal FAT10 degradation depends on Nub1

To elucidate how Nub1 mediates FAT10 degradation, we reconstituted this process *in vitro* with *E. coli* expressed full-length human FAT10 and Nub1, as well as human 26S (hs26S) proteasome isolated from HEK293 cells (Figure 1A). Using SDS-PAGE, we monitored the degradation of FAT10 in the absence or presence of excess Nub1, and found that, at least *in vitro*, FAT10’s rapid turnover strictly depends on Nub1 and does not require ubiquitination (Fig. 1B). Control experiments with the proteasome-specific inhibitor MG132, the slowly hydrolyzed ATP analog ATPγS, or the Rpn11 inhibitor ortho-Phenanthroline (oPA) confirmed that this degradation relies on proteolysis by the 20S core peptidase as well as the ATP-dependent unfolding and translocation by the proteasomal 19S RP, yet is independent of Rpn11-mediated deubiquitination (Figure 1C).

**Figure 1:**
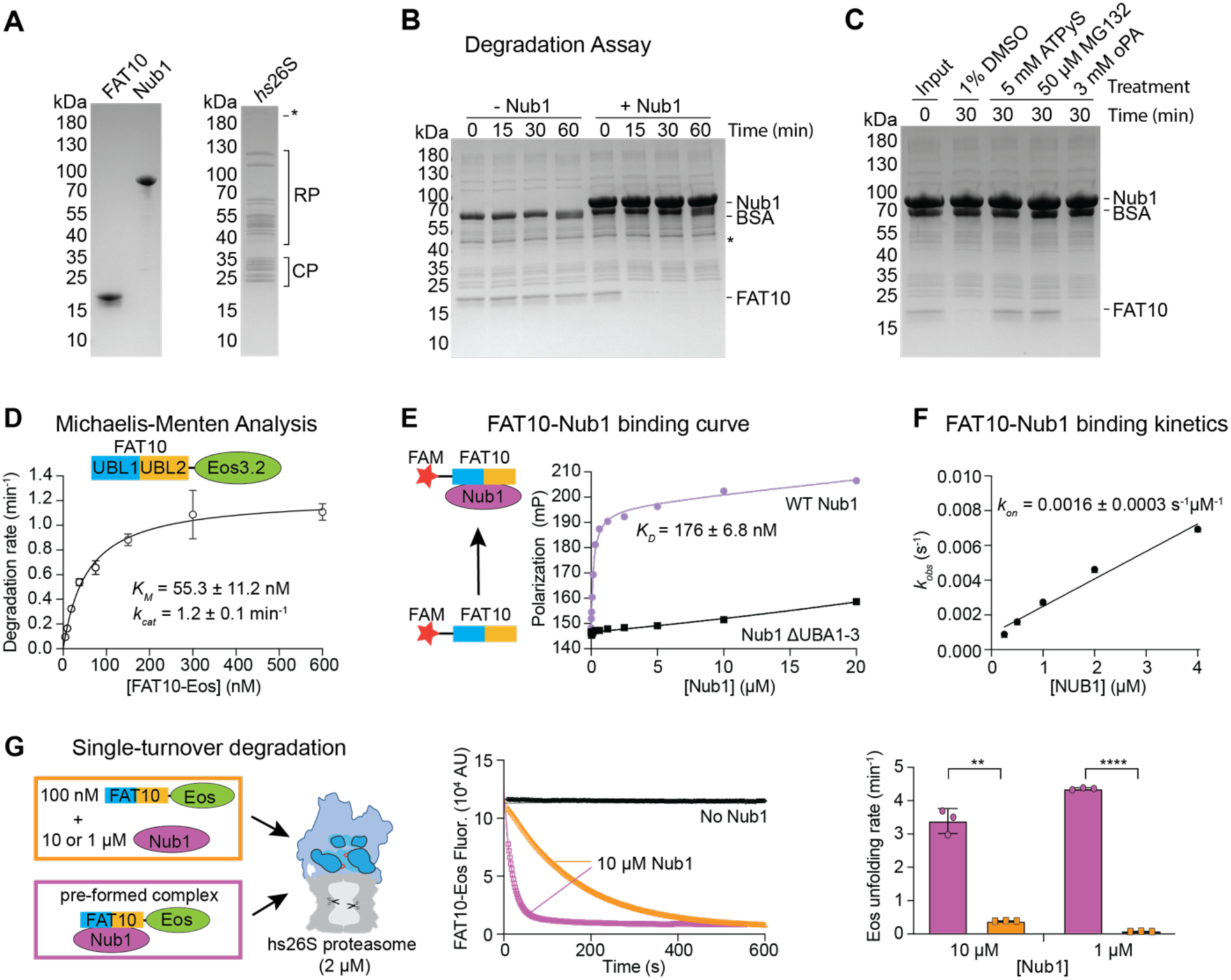
Ubiquitin-independent FAT10 degradation by the *hs*26S proteasome depends on Nub1 complex formation as the rate-limiting step in degradation. **A**) Coomassie-stained SDS-PAGE gels documenting the purity of recombinantly expressed human FAT10 (3 μg) and human Nub1 (1.5 μg), and endogenous *hs*26S proteasome purified from HEK293 cells (1.5 μg). **B)** FAT10 degradation by the *hs*26S proteasome depends on the presence of Nub1. SDS-PAGE analysis of aliquots taken at different times during the degradation of FAT10 (5 μM) by the *hs*26S proteasome (100 nM) in the absence or presence of Nub1 (15 μM). **C)** SDS-PAGE analysis of FAT10 degradation by *hs*26S proteasome in the presence of indicated concentrations of DMSO, the non-hydrolyzable ATP analog ATPγS, the proteasome inhibitor MG132, and the Rpn11 deubiquitinase inhibitor o-phenanthroline (oPA). **D)** Michalis-Menten analysis of FAT10-Eos3.2 degradation (10 – 600 nM) by *hs*26S proteasome (2 nM) in the presence of excess Nub1 (10 μM). Shown are the mean values and standard deviations of the degradation velocity determined from the loss of Eos fluorescence for n = 3 technical replicates. **E)** Measurement of Nub1-FAT10 complex formation by fluorescence polarization. ^FAM^FAT10 (100 nM) was incubated with varying Nub1 concentrations for 45 mins on ice before measuring the polarization. A truncated Nub1^ΔUBA1–3^ variant was used as a control for non-specific binding. Data shows mean +/− SD (n=3) **F)** Kinetic analysis of the slow complex formation between FAT10 and Nub1. Shown are the apparent rate constants *k_obs_* for the binding of ^FAM^FAT10 (20 nM) to Nub1 at varying concentrations (250 – 4000 nM) as determined from the change in fluorescence polarization. Data shows mean +/− SD (n=3) **G)** Complex formation of Nub1 and FAT10 is the rate limiting step in FAT10 degradation. Left: Schematic for the experimental setups of the single-turnover degradation reactions, in which *hs*26S proteasome (2 μM) was mixed with either the individual components (orange outline) of FAT10-Eos (100 nM) and excess Nub1 (1 or 10 μM) or preformed complexes (purple outline) of FAT10-Eos and Nub1 at the same concentrations. Middle: Representative curves for the loss of FAT10-Eos fluorescence during the single-turnover degradation in the absence of Nub1 (black) or the presence of Nub1 (10 μM) without (orange) or with (purple) preincubation for 30 min. Right: Shown are the mean values and standard deviations for the FAT10-Eos3.2 unfolding by the *hs*26S proteasome in the presence of Nub1 (1 or 10 μM) with (purple) or without pre-incubation for Nub1/FAT10 complex formation. N = 3 technical replicates. Statistical significance was calculated using an unpaired two-tailed Welch’s t test: ∗∗∗∗p < 0.0001; ∗p = 0.0446.

Interestingly, it was previously shown that overexpressed FAT10 is degraded by the 26S proteasome in yeast cells that naturally lack Nub1, and co-expressing Nub1 accelerated this *in vivo* degradation ^36^. However, when reconstituting this process *in vitro*, we found that the 26S proteasome from yeast *S. cerevisiae* (*sc*26S), similar to its human counterpart, cannot significantly degrade FAT10 in a Nub1-independent manner (Figure S1A). Hence, other mechanisms may aid FAT10 degradation in yeast cells, for instance ubiquitination and/or Cdc48-mediated unfolding. FAT10 was previously reported to easily aggregate ^34,44^ and be susceptible to degradation by the isolated 20S CP after longer incubations *in vitro* ^45^. However, our *E.coli*-expressed wild-type FAT10 was highly soluble, well-behaved, and not truncated during expression or purification (Figure S1B). It was only very slowly degraded by the yeast 20S CP alone (Figure S1C) or by the *sc*26S proteasome in the presence of ATPγS (Figure S1D), likely due to its intrinsic lability and some extent of spontaneous unfolding over the 60 min period of the experiment. These observations support the validity of our findings that FAT10 is rapidly and specifically degraded by human and yeast 26S proteasomes in a process that strongly depends on ATP and Nub1.

### Nub1 targets FAT10 conjugates for degradation

To quantitatively measure the kinetics of FAT10 turnover by the *hs*26S proteasome, we designed a reporter construct with FAT10 fused to the N-terminus of monomeric mEos3.2 (FAT10-Eos), which allowed monitoring of substrate unfolding and degradation through the loss Eos fluorescence. mEos3.2 is a well-folded protein that lacks unstructured initiation regions suited for proteasomal engagement, such that even in its ubiquitinated form it is not degraded, but requires prior unfolding by Cdc48 ^5^. However, we found that FAT10-Eos is robustly degraded by the *hs*26S proteasome in the presence of Nub1 (Figure S1E). As expected for a fusion with the hard-to-unfold Eos domain, the observed rate was lower than for the isolated FAT10 (Figure 1B, S1E). Michaelis-Menten analysis in which we titrated FAT10-Eos in the presence of saturating Nub1 concentrations revealed a *K_M_* of ∼ 55.3 ± 11.2 nM and a *k_cat_* of ∼ 1.2 ± 0.1 min^−1^ (Figure 1D), in good agreement with our previously reported velocity for the degradation of ubiquitinated mEos3.2 by the *sc*26S proteasome ^4^.

Given that FAT10 may function as a targeting signal as well as an initiation region for proteasomal degradation, we wondered whether adding a long-disordered tail to the C-terminus of FAT10-Eos was enough to bypass the Nub1 requirement and allow engagement by the proteasome, similar to our previous findings for the Cdc48 dependence of untailed versus tailed ubiquitinated mEos3.2 for proteasomal degradation ^5^. However, we could not detect degradation of FAT10-Eos-tail by the *hs*26S proteasome in the absence of Nub1 (Figure S1F), indicating that FAT10 alone is insufficient for either recruitment or initiation. To explore this further, we created a linear fusion of four ubiquitin moieties with Eos-tail (Ub_4_-Eos-tail), which, in contrast to the untailed Ub_4_-Eos control, was degraded by the *hs*26S proteasome, albeit slowly (Figure S1G). These results indicate that, in the absence of Nub1, FAT10 either does not interact with the human proteasome or binds in a way that is not compatible with presenting the C-terminal tail on Eos for proteasomal engagement. Nub1 has previously also been identified to accelerate the degradation of the NEDD8 ubiquitin-like modifier, and we therefore tested the degradation of NEDD8-Eos and NEDD8-Eos-tail fusions in the absence and presence of Nub1. Neither construct showed a significant turnover (Figure S1H), suggesting that there is a specific FAT10-Nub1 interaction driving the degradation of FAT10-ylated proteins by the *hs*26S proteasome.

Interestingly, FAT10-Eos-tail and NEDD8-Eos-tail were degraded by the *sc*26S proteasome in the absence of Nub1 (Figure S1I,J), indicating that both modifiers are sufficient to deliver a substrate for degradation, provided that a long-disordered tail for engagement is present on the substrate. The yeast proteasome thus appears more promiscuous in UBL-domain binding, and indeed previous work showed that many types of UBL-fused substrates can be degraded by the *sc*26S proteasome ^46–49^. Importantly, FAT10 is not sufficient to mediate the degradation of tailless FAT10-Eos by the *sc*26S proteasomes, and still depends on Nub1 (Figure S1I). For the degradation by *hs*26S proteasome, we can therefore conclude that Nub1 is required for both the specific recruitment and the initiation of the FAT10-ylated substrate.

### FAT10 and Nub1 slowly form a high-affinity complex

For investigating the mechanisms by which Nub1 enables FAT10 degradation, we first used size-exclusion chromatography and observed that FAT10 and Nub1 form a stable 1:1 complex (Figure S2A). To determine their affinity in fluorescence-polarization-based binding measurements, we attached a fluoresceine-amidite (FAM)-modified peptide through sortase labeling to the N-terminus of FAT10. When mixing this ^FAM^FAT10 with excess Nub1, we detected a slow increase in polarization that was Nub1-concentration dependent (Figure 1E,F, Figure S2B). Titrating Nub1 and analyzing the polarization endpoints revealed a *K_D_* of 176.0 ± 6.8 nM for the Nub1/ ^FAM^FAT10 complex (Figure 1E). This represents an approximate dissociation constant, as some aggregation occurred at higher Nub1 concentrations, potentially caused by either the nature of the Nub1/FAT10 interaction or the hydrophobic FAM label on FAT10. Deleting Nub1’s UBA domains eliminated FAT10 binding (Figure 1E), which agrees with previous reports ^41^ and confirms a specific interaction between Nub1 and FAT10. Measuring the kinetics revealed an association constant of *k_on_* = 0.0016 ± 0.003 s^−1^μM^−1^ (Figure 1F).

To further explore the importance of this slow but tight complex formation, we performed Nub1-mediated FAT10-Eos degradation experiments under single-turnover conditions, i.e. in the presence of excess *hs*26S proteasome, which provides insight into to processes prior to mEos3.2 unfolding. When mixing FAT10-Eos (100 nM) with saturating amounts of Nub1 (10 µM) and *hs*26 proteasome (2 µM, see Figure 1D), we observed a single-exponential decay of Eos fluorescence with a time constant of *τ* = 162 ± 4.3 s, equivalent to a degradation rate of *k_unfold_* = 0.37 min^−1^ (Figure 1G). This represents the time required for Nub1/FAT10-Eos complex formation, binding to the proteasome, unfolding and translocation of FAT10, and initial unraveling of the Eos β-barrel. In contrast, when we performed the experiment at identical concentrations, but pre-incubated FAT10-Eos and Nub1 for 20 min prior to *hs*26 proteasome addition, we detected fast processing with a time constant of *τ_fast_* ∼ 18.0 ± 2.2 s. We also observed a low-amplitude (9%) second phase with a time constant of *τ_slow_* = 192 ± 31.7 s (Figure 1H), which we attribute to malformed or aggregated Nub1/FAT10-Eos complex that was present in small amounts after the pre-incubation. Importantly, the dominant first phase of the degradation reaction proceeded almost an order of magnitude faster than the unfolding observed without pre-incubating Nub1 and FAT10-Eos (*k_unfold_^(pre-inc.)^* = 3.3 min^−1^ versus *k_unfold_* = 0.37 min^−1^; Figure 1G). We validated this by reducing the Nub1 concentration to 1 µM, which led to even slower degradation kinetics when not pre-incubating Nub1 and FAT10-Eos (Figure 1G), as their complex formation is rate-determining and concentration dependent. In contrast, degradation progressed still rapidly when using the pre-formed Nub1/FAT10-Eos complex at identical concentrations of 100 nM FAT10 and 1 µM Nub1 (Figure 1H). Together, the polarization-based binding measurements and the degradation studies under single-turnover conditions demonstrate that the complex formation between Nub1 and FAT10 is slow and rate-determining for proteasomal turnover.

### Nub1 traps partially unfolded FAT10 by binding to a single beta strand

Because the 26S proteasome requires an unstructured initiation region to engage a substrate for degradation, we wondered whether Nub1’s binding to FAT10 played any role in providing a flexible segment and how the two proteins interact. To assess changes in the conformation and solvent accessibility of FAT10 upon binding to Nub1, we employed hydrogen-deuterium exchange monitored by mass spectrometry (HDX-MS). Protonated FAT10 in the absence or presence of excess Nub1 was incubated in D_2_O for variable times before quenching, pepsin digest, and peptide detection by LC/MS. Under both conditions we observed excellent peptide coverage spanning the entire FAT10 sequence (Figure 2A, Table S1). Interestingly, several peptides from both UBL domains of FAT10 exhibited bimodal distributions in the absence of Nub1, with a rapidly exchanging population already detected at the earliest time point (Figure 2B,C; Figure S3). This indicates the presence of exposed, i.e. unfolded or partially unfolded states, in addition to a protected folded state, and it is consistent with FAT10’s dynamic nature previously suggested based on *in vivo* degradation, molecular dynamics, and biophysical measurements ^34^. The exchange of some peptides appears to show a mixture of EX1 and EX2 kinetics, which, together with overlapping peaks in the mass spectra, made it difficult to fit bimodal distributions and determine the relative populations for each state (Figure S3A). We therefore used the left peak from bimodal distributions to compare the differences between free FAT10 and the FAT10/Nub1 complex (Figure 2A). Except for UBL1’s last beta strand, which showed protection in the presence of Nub1, Nub1 binding caused a strong exposure of peptides spanning the entire UBL1 domain (Figure 2A), suggesting that it induces or traps an unfolded state of FAT10’s UBL1. We also focused on the presence or absence of bimodal distributions to describe the effect of Nub1 binding on FAT10, and selected four example peptides, two from each UBL domain. Three of the peptides displayed clear bimodal deuterium uptake in the absence of Nub1, with a slowly exchanging and a fully exchanged population throughout all early time points, whereas the fourth peptide, derived from UBL1’s last beta strand, showed primarily unimodal distribution (Figure 2B,C). Remarkably, Nub1 binding eliminated the bimodal distribution for the first UBL1 peptide, leaving only the fully exposed population, whereas both UBL2 peptides stayed unaffected and retained bimodal exchange (Figure 2B,C; Figure S3A,B). This indicates that Nub1 specifically interacts with UBL1 and has no effect on UBL2, which would be consistent with previous studies indicating that FAT10’s UBL1 and UBL2 represent independently folding domains with no considerable interactions ^34^. Interestingly, the slowly exchanging UBL1 population in the absence of Nub1 shows deuterium-uptake kinetics in the minute range, similar to the time constants we observed for the Nub1/FAT10 complex formation in our fluorescence-polarization and single-turnover degradation experiments (Figures 1F,G). We therefore propose that Nub1 uses conformational selection to bind and trap the spontaneously unfolding UBL1 domain of FAT10, rather than actively inducing its unfolding. UBL1’s last beta strand, which shows protection from deuterium exchange upon Nub1 binding, follows H75 and we therefore term it H75^beta-strand^ (Figure 2B,C). This single beta strand appears to be Nub1’s binding site within FAT10.

**Figure 2:**
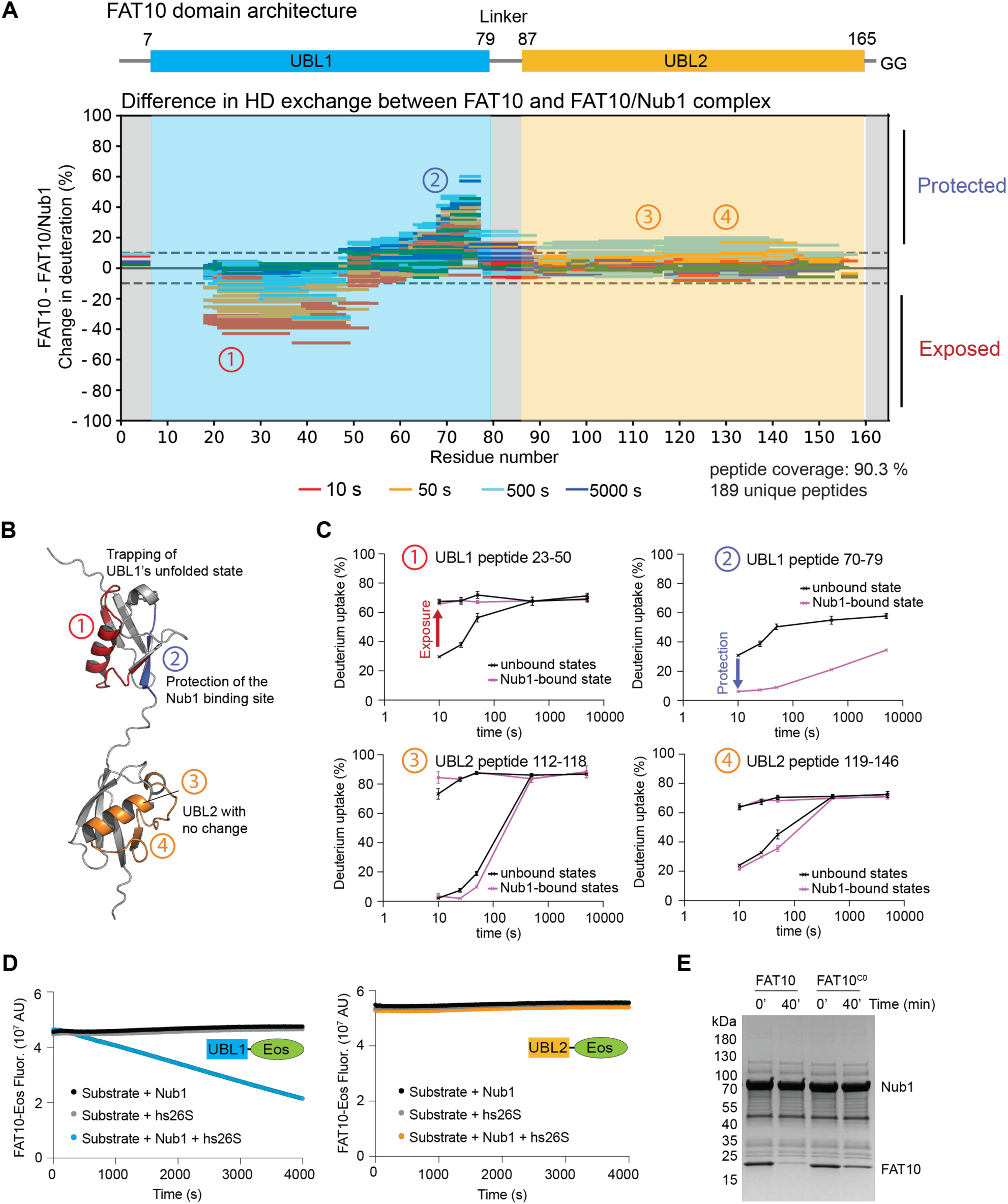
Nub1 stabilizes the unfolded state of FAT10’s UBL1. **A**) Top: schematic of FAT10’s domain architecture, with the UBL1 domain shown in cyan and the UBL2 domain in orange. Bottom: Wood’s plot representation of the percent changes in deuteration between free FAT10 and the FAT10/Nub1 complex, with decreased uptake (protection) shown above 0% and increased uptake (exposure) upon complex formation shown below 0%. For bimodal peptides only less exchanged peaks are shown for comparison. The time of HD exchange (10 – 5000 s) is indicated by different colors. Changes in deuteration between the dotted lines were considered to be not significant. Encircled numbers indicate four representative peptides in the UBL1 and UBL2 domains that get exposed (red), protected (blue), or show no change upon FAT10/Nub1-complex formation. Their positions within the FAT10 structural model are shown in B), and their deuterium uptake kinetics are depicted in C). **B)** Structure representation of FAT10, indicating the positions of peptide 1 (red) that shows increased exposure upon complex formation with Nub1, peptide 2 (blue) that represents the last beta strand in the UBL1 domain and gets protected by Nub1 binding, and peptides 3 and 4 (orange) in the UBL2 domain that are unaffected by Nub1 binding. **C)** Deuterium-uptake plots for peptides 1 – 4 depicted in panels A) and B). Shown are the means and standard deviations for the percentages of deuterium uptakes based on the theoretical maximum deuteration for each peptide and determined from N = 3 technical replicates. Top left: peptide 1 shows a bimodal distribution of deuterium uptake in the absence of Nub1 and becomes unimodally exposed throughout the time course upon FAT10 binding. Top right: peptide 2 shows unimodal exchange and becomes protected upon FAT10 binding. Bottom left and right: the bimodal uptake behavior for peptides 3 and 4 throughout the time course remains unchanged after FAT10 binding Nub1. **D)** Example Eos-fluorescence traces for degradation of FAT10^ΔUBL2^-Eos (left) and FAT10^ΔUBL1^-Eos (right) by hs26S proteasome with (blue and orange) or without (grey) excess of Nub1 (15 μM). **E)** Coomassie-stained SDS-PAGE gel showing the end points for the multiple-turnover degradation of wild-type FAT10 and the thermodynamically stabilized cysteine-free FAT10^C0^ by hs26S proteasome after pre-incubation with excess Nub1 (15 μM).

### Nub1 delivers FAT10’s unfolded UBL1 domain for proteasomal engagement

Since FAT10’s UBL1 domain seems to provide both the binding site for Nub1 and the disordered initiation region for engagement by the proteasome, we tested whether the presence of this domain is sufficient to facilitate the degradation of mEos3.2. Indeed, our single-turnover experiments showed that FAT10^ΔUBL2^-Eos is degraded in a Nub1-dependent manner, albeit ∼ 2-fold more slowly than full-length FAT10 (*τ ^FAT^*^10^ *^ΔUBL^*^2^ = 54.4 +/− 4.5 s vs. *τ ^FAT^*^10^ *^WT^*= 28.6 +/− 0.9 s; Figure S4A). These findings indicate a kinetic effect of FAT10’s UBL2 domain on the rate-limiting step during initiation rather than a contribution to the binding affinity, and this domain may form minor interactions that help orient the Nub1/FAT10 complex for initiation or act as a spacer between the Nub1-bound UBL1 domain and the protein substrate to prevent steric clashes with the proteasome. In contrast, FAT10^ΔUBL1^-Eos showed no degradation (Figure 2D), even after attaching an unstructured segment to the C-terminus (FAT10 ^ΔUBL1^-Eos-tail, Figure S4B). It was previously proposed that FAT10’s degradation initiates on its N-terminus ^34^, and to prove this in our reconstituted system, we blocked the N-terminus with a fusion to Smt3, the yeast homolog of the Small Ubiquitin-like Modifier, SUMO. No degradation was observed for this Smt3-FAT10 construct (Figure S4C), confirming that a free N-terminus is critical for FAT10 degradation. Based on our HDX-MS experiments and the slow Nub1/FAT10 complex formation, we hypothesized that the intrinsic lability and spontaneous unfolding of FAT10’s UBL1 domain are critical for Nub1 binding and consequently degradation by the proteasome. Previous studies showed that a quadruple Cys-to-Ala mutant, FAT10^C0^, with increased thermodynamic stability exhibited decreased degradation *in vivo* ^34^. We therefore generated FAT10^C0^ and, indeed, observed that degradation by the *hs*26S proteasome was strongly decelerated (Figure 2E) and Nub1 binding was undetectable by fluorescence polarization (Figure S4D). Together, these results demonstrate that FAT10’s UBL1 domain functions as a degradation-initiation region whose structural instability allows Nub1 to bind and trap the unfolded state for engagement by the 26S proteasome.

### Nub1 domains form an expandable channel for FAT10 binding

Based on previously annotated domains ^41^ and secondary structure predictions ^50^, Nub1 contains a N-terminal domain (NTD) followed by an UBL domain that is attached through helical and unstructured linkers to a core domain. This core fold leads into the UBA1 domain, which is connected through another helical linker to the UBA2 and UBA3 domains, followed by a long-disordered region and two C-terminal helices. To assess changes in the conformation and solvent accessibility of Nub1 upon FAT10 binding, we performed HDX-MS experiments with protonated Nub1 in the absence or presence of excess FAT10. FAT10 binding led to changes in hydrogen exchange across the entire sequence of Nub1, and the differences in the exchange for peptides from the unbound and bound samples at various time points are shown in Figure 3A (see also Table S1). Slowly exchanging peptides for unbound and FAT10-bound Nub1 exhibit a good correlation with the folded domains, whereas fast-exchanging peptides match with predicted linker regions (Figure 3A, Figure S5). The observed differences in the exchange profiles for unbound versus FAT10-bound Nub1 indicate considerable conformational changes, including an exposure of peptides in Nub1’s NTD and UBL domains, and deuterium uptake plots for several selected Nub1 peptides in the absence and presence of FAT10 are shown in Figure S6.

**Figure 3:**
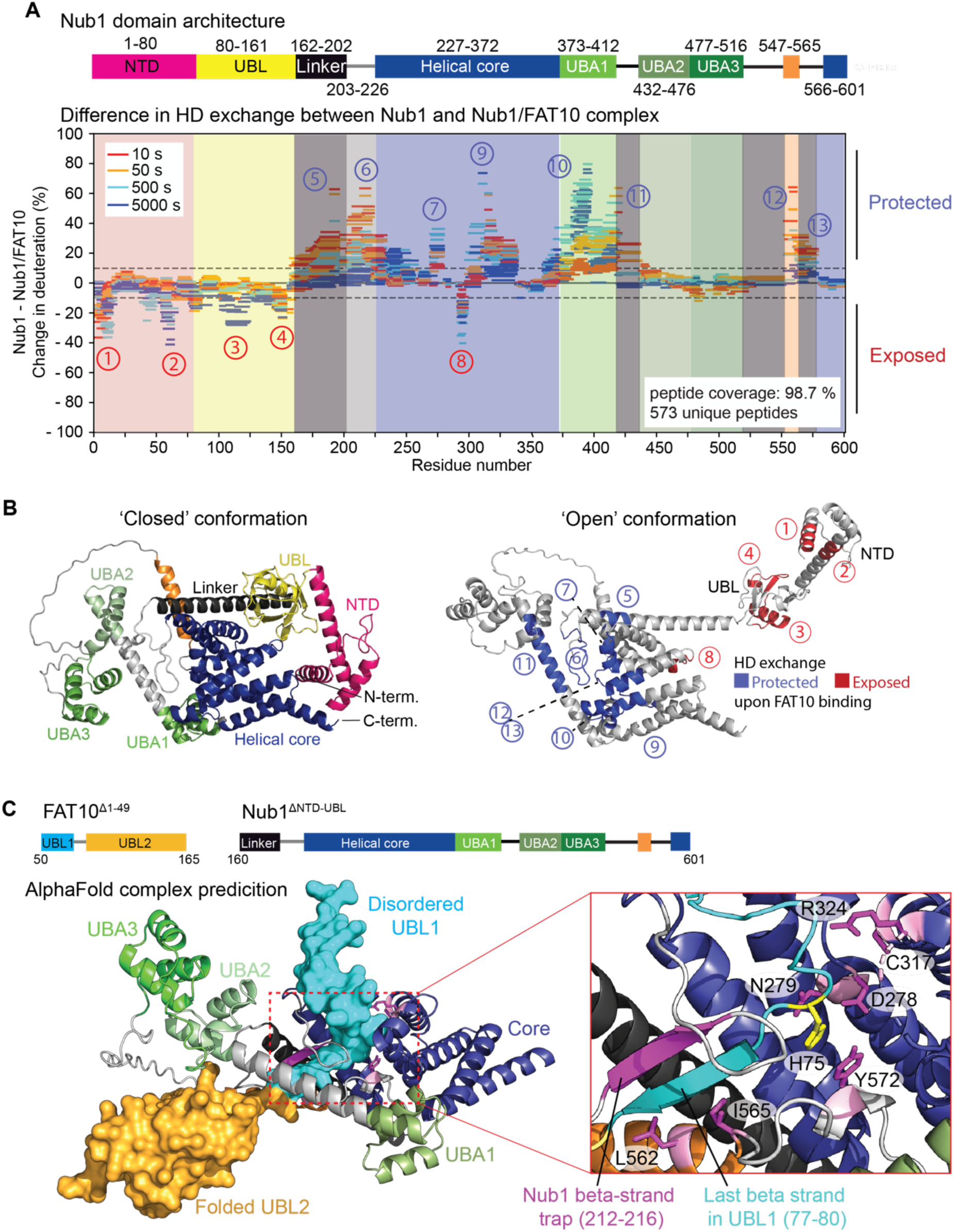
Hydrogen-deuterium exchange reveals conformational changes in Nub1 upon trapping FAT10 in a partially unfolded state. **A**) Top: schematic of Nub1’s domain architecture and with residue numbers indicated. Bottom: Wood’s plot representation of the percent changes in deuteration between free Nub1 and the Nub1/Fat10 complex, with decreased deuterium uptake (protection) upon FAT10-complex formation for individual peptides shown above 0% and increased uptake (exposure) upon complex formation shown below 0%. The time of HD exchange (10 – 5000 s) is indicated by different colors. Changes in deuteration between the dotted lines were considered to be not significant. Encircled numbers indicate specific peptides that get exposed (red) or protected (blue) upon FAT10-complex formation, and their position within the Nub1 structural model is shown in B). **B)** CollabFold-generated AlphaFold models of the Nub1 structure show the domain architecture and distinct conformational states. Assigned domains and their boundaries based on AlphaFold models are indicated by colors and consistent with the domain predictions shown in A). The positions of UBA2, UBA3, NTD, and UBL domains vary between different models, in which the UBL domain is observed docked against the core domain in a ‘closed’ Nub1 conformation (left) or exposed in an ‘open’ Nub1 conformation (right). Selected peptides highlight key changes in the open-state Nub1 upon FAT10 binding (right), with more protected peptides shown in blue and more exposed peptides in red mapped onto the Nub1 structure. Peptides with increased protection in the FAT10-bound complex are generally localized within areas and linkers that line a channel formed between the helical core domain and the UBA domains of Nub1. **C)** AlphaFold Multimer structure prediction of the Nub1/FAT10 complex. Top: schematics for the N-terminally truncated variants of Nub1 and FAT10 that in addition to the full-length versions (see Fig. S9) were used for the structure prediction of the Nub1/FAT10 complex by AlphaFold Multimer. Bottom left: Structural model for the Nub1/FAT10 complex, with Nub1 shown in ribbon representation, colored as in the schematic at the top, and FAT10 depicted in surface representation, with the UBL1 domain threaded through Nub1 and trapped in an unfolded state. Bottom right: Zoom-in view of the Nub1-FAT10 interaction, where the last beta strand of FAT10’s UBL1 domain (cyan ribbon) is trapped by forming an anti-parallel beta sheet with the “beta-strand” (bs) linker (purple ribbon) of Nub1. FAT10’s H75 (yellow stick representation) is coordinated by D278, N279 and Y572 of Nub1, and additional Nub1 residues relevant for the interaction with FAT10 are shown in purple stick representation.

AlphaFold structure predictions revealed at least three conformations for the isolated Nub1 that we term ‘closed’, ‘open’, and ‘partially open’, based on the position of Nub1’s UBL domain (Figure 3B, Figure S7A,B). Together, the core domain, the three UBA domains, and their linkers form a loop structure that is anchored through extensive interactions between the C-terminal helices and the core. This topology explains why we observed low solubility for truncated Nub1 that lacks the C-terminal helices, whereas variants with deleted NTD-UBL (Nub1^ΔNTD-UBL^ and Nub1^ΔNTD-UBL-Linker^) or deleted UBA domains (Nub1^ΔUBA1–3^) are well behaved and most likely properly folded. There are consequently three segments to Nub1: the NTD-UBL, the core domain, and the looped-out region containing UBA1, UBA2, and UBA3. It appears that the flexible linker between UBA3 and the C-terminal helices allows this looped region to open and close.

Notably, in our HDX-MS experiments, many regions surrounding and lining the channel formed between Nub1’s core body and the three UBA domains become protected upon FAT10 binding, including a flexible linker within the channel that is well conserved throughout evolution (Figure 3A,B, Figure S8). This protection pattern suggests that FAT10 somehow interacts with this linker and the channel through Nub1. Furthermore, FAT10 binding leads to an increased exposure of Nub1’s UBL domain and a region on Nub1’s core that the UBL domain contacts in the closed conformation, indicating that FAT10 induces an open Nub1 state, with the UBL domain exposing an interface that is equivalent to ubiquitin’s I44 patch. The I44 patch of ubiquitin is typically involved in binding various interaction partners, including the ubiquitin receptors of the proteasome. We therefore postulate that free Nub1 primarily resides in a closed state, while FAT10 binding induces the open conformation with an exposed UBL domain that interacts with a proteasomal receptor.

### Structural model for Nub1 with trapped FAT10

To gain further insight into the Nub1/FAT10 complex and corroborate our HDX-MS-based models, we used structure predictions with AlphaFold-Multimer ^51^ for either truncated sequences of FAT10 (residues 50-166) and Nub1 (residues 161-600) (Figure 3C, S7C) or their full-length versions (Figure S9). Remarkably, we were able to obtain several confident models in which FAT10’s H75^beta-strand^ is inserted into a channel between Nub1’s helical core and UBA domains, while other parts of FAT10’s UBL1 domain are disordered (Figure 3C). All predicted interactions between Nub1 and FAT10 are consistent with our HDX-MS analyses (Figures 2A and 3A), including multiple regions throughout Nub1 that surround FAT10’s H75^beta-strand^ and showed protection from deuterium exchange upon FAT10 binding. The H75^beta-strand^ forms an anti-parallel beta-sheet with the region in Nub1 that represents the unstructured linker between the UBL and core domains in the absence of FAT10 but exhibits strong protection in HDX-MS upon FAT10 binding, and we refer to this as the beta-strand trap (bs-trap) (Figure 2C, Figure 3C). In addition to the backbone interactions between these beta strands, H75 of FAT10 is coordinated by multiple conserved Nub1 residues: D278, N279 and Y572 (Figure 3C), which all get protected from deuterium exchange in the presence of FAT10. As most of the contacts in the antiparallel beta sheet between bs-trap and H75^beta-strand^ are mediated by the peptide backbones, we decided to disrupt these interactions by deleting part of the bs-trap linker or by mutating two residues to proline. Both Nub1^Δbs-trap^ and Nub1^D214P/A216P^ abolished degradation of FAT10-Eos (Figure 4A) and showed no binding to ^FAM^FAT10 (Figure 4B). Furthermore, alanine mutations of several Nub1 residues that in the AlphaFold-Multimer models coordinate FAT10’s H75^beta-strand^ throughout Nub1’s binding channel led to compromised FAT10-Eos degradation and ^FAM^FAT10 binding (Figures 4A,B, Figure S10A-B).

**Figure 4:**
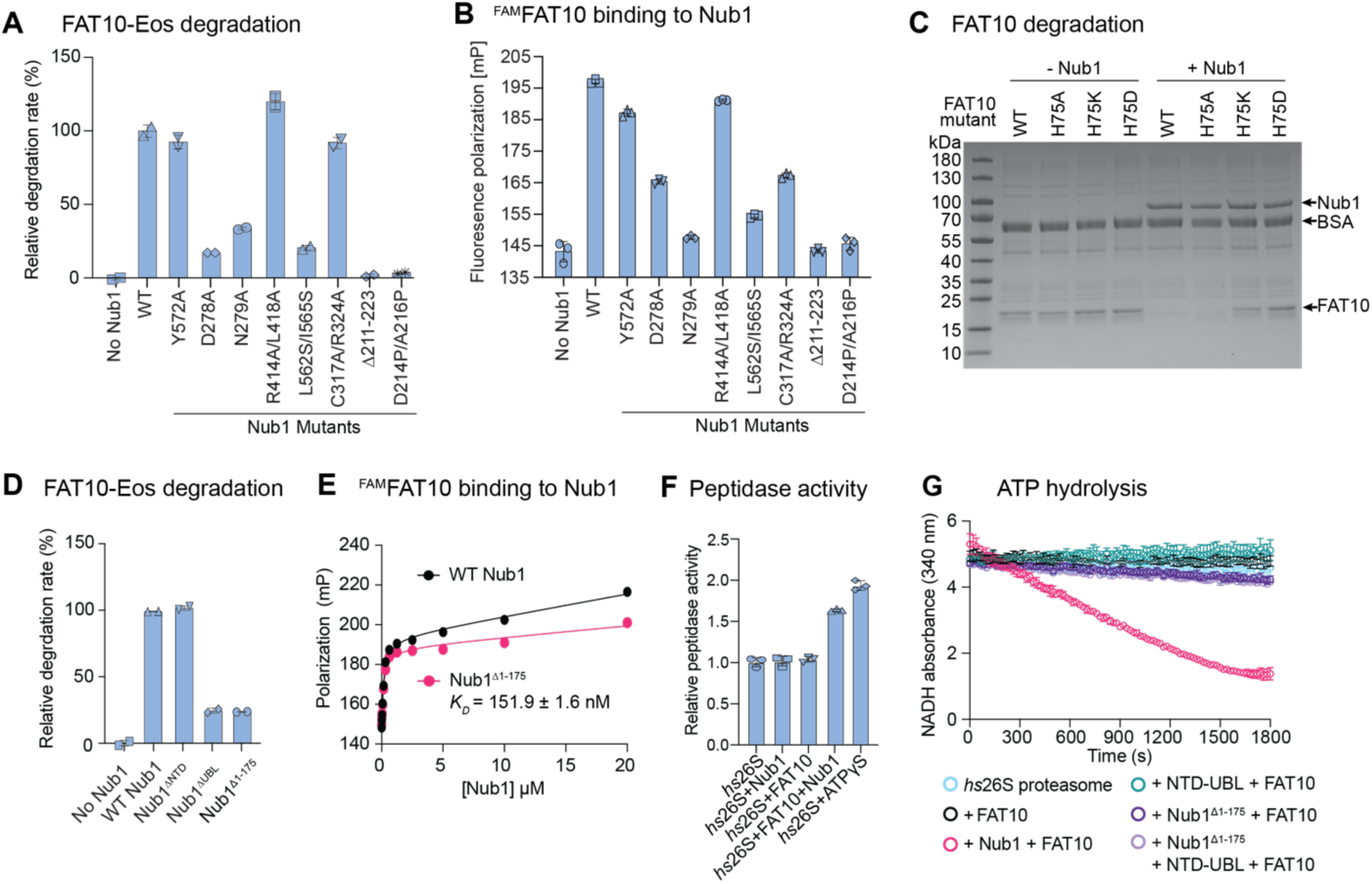
Nub1 uses a beta-strand trap to bind FAT10 and its UBL domain for interaction with the proteasome. **A**) Relative rates for the multiple-turnover degradation of FAT10-Eos (1 μM) by the *hs*26S proteasome (100 nM) in the presence of wild-type Nub1 (set to 100 %) or Nub1 mutants (5 μM). Shown are the mean values and standard deviation for N = 3 technical replicates. **B)** Fluorescence polarization as a readout for the binding of ^FAM^FAT10 (100 nM) to wild-type Nub1 and its mutants (5μM) after pre-incubation for 30 min. **C)** Coomassie-stained SDS-PAGE gel showing the endpoints for the degradation of FAT10 wild-type and its H75 mutants (10μM) by *hs*26S proteasome (100 nM) in the absence and presence of Nub1 (2.5 μM) after 30 minutes. **D)** Relative rate of FAT10-Eos degradation (1 μM) by hs26S proteasome (100 nM) in the presence of various Nub1 truncation mutants (5 μM). Bar graphs show the mean values and error bars represent the standard deviation for N = 3 technical replicates and normalized to the degradation in the presence of wild-type Nub1. **E)** Fluorescence-polarization measurement of the Nub1-FAT10 complex formation indicates that Nub1’s UBL domain is dispensable for FAT10 binding. ^FAM^FAT10 (100 nM) was incubated with varying concentrations of wild-type Nub1 or Nub1^Δ1–175^ for 45 mins on ice before measuring the polarization. Data shows mean +/− SD (N = 3). **F)** Peptidase activity of the *hs*26S proteasome in the presence of FAT10, Nub1, the FAT10/Nub1 complex, or ATPγS instead of ATP. Shown are the mean values and standard deviations of the peptidase activities for N = 3 technical replicates, normalized to the activity of *hs*26S proteasome in the presence of ATP. **G)** Example traces for the depletion of NADH in a coupled ATP-hydrolysis assay with *hs*26S proteasome in the absence or presence of FAT10, Nub1, Nub1^Δ1–175^, or a NTD-UBL fragment of Nub1.

In additions to Nub1 regions that are in direct contact with FAT10, we observed protection from deuterium exchange upon FAT10 binding for a helix in the UBA1 domain that is positioned right behind Y572 in Nub1 (Figure S10C). The UBA1 domain may thus dock against and stabilize the segment around Y572 that coordinates the linchpin residue H75 in FAT10, whereas in the absence of FAT10 this domain appears more mobile, potentially facilitating FAT10’s insertion and trapping within the Nub1 channel. Indeed, deletion of Nub1’s UBA1 domain is sufficient to inhibit FAT10 degradation, whereas the deletion of UBA2 and UBA3 has no major effects (Figure S10D). Finally, we made three mutations of FAT10’s critical H75 residue (H75A, H75D, and H75K) and tested their influence on degradation and Nub1 binding. All three mutants showed compromised binding to Nub1 (Figure S4D) and inhibited degradation (Figure 4C), with H75D and H75K completely abolishing any turnover. Our findings from mutational and HDX-MS studies are thus in excellent agreement with the AlphaFold-Multimer models, which provide the first structural picture of how Nub1 traps unfolded FAT10. Based on these results, we propose Nub1 to be an ATP-independent chaperone that traps the last beta strand in FAT10’s UBL1 domain and thereby stabilizes the unfolded state for proteasomal engagement and degradation.

### Nub1’s UBL domain is critical for FAT10 delivery to the proteasome

Based on the results presented above, we predicted that FAT10 binding induces the open state of Nub1 and allows the UBL domain to contact a proteasomal receptor. Through this mechanism, uncomplexed Nub1 would be in a closed state and not compete with FAT10-bound Nub1 for proteasome binding. To test this, we made mutants of Nub1 with either the NTD, UBL, or both domains deleted (Nub1^ΔNTD^, Nub1^ΔUBL^, and Nub1^Δ1–175^) and analyzed their activity in facilitating FAT10-Eos degradation. While the NTD deletion had no effect, eliminating the UBL domain strongly compromised FAT10 degradation (Figure 4D, Figure S11A). As expected, the binding of ^FAM^FAT10 was not affected by the deletion of the NTD and UBL domains in Nub1^Δ1–175^ (Figure 4E). Nub1’s UBL domain is thus dispensable for FAT10 binding, but critical for robust FAT10 degradation by the 26S proteasome. Consistent with our model that free Nub1 primarily exists in a closed conformation with its UBL tucked-in and therefore unable to contact the proteasome, we did not observe inhibition of Nub1-mediated FAT10 degradation when higher concentrations of excess, free Nub1 were present (Figure 1B,D,G).

It was previously reported that UBL domains, including the one of Nub1, allosterically activate the proteasomal peptidase or ATPase activities ^42,52^. When testing the peptidase-activity response of *hs*26S proteasome, we observed a stimulation only after adding FAT10 and Nub1 together, while Nub1 and FAT10 individually had no effect (Figure 4F). This may indicate that the peptidase stimulation either requires a FAT10-bound, open conformation of Nub1 with exposed UBL domain or that it is caused by substrate engagement and degradation, which shifts the proteasome from non-processing to processing states with an open 20S peptidase gate. To explore this further, we measured the proteasomal ATPase activity in the absence and presence of FAT10, Nub1, or FAT10+Nub1 (Figure 4G). The *hs*26S proteasome by itself had undetectable ATPase activity, which may be a consequence of it residing primarily in the non-processing conformational state, as judged by cryo-EM particle distributions ^53^. Again, robust stimulation was only observed when adding both Nub1 and FAT10 (Figure 4G). Importantly, a NTD-UBL fragment of Nub1 caused no ATPase stimulation, indicating that UBL-domain binding has no allosteric effects on the proteasomal activities, and the observed stimulation by Nub1 and FAT10 is likely due to active degradation and the conformational shift to substrate-processing states with increased ATP hydrolysis and an open gate of the 20S CP. Furthermore, adding the NTD-UBL fragment together with the complementary N-terminal deletion variant Nub1^Δ1–175^ in complex with FAT10 also did not stimulate the proteasomal ATPase activity (Figure 4G), which rules out that the UBL domain activates the proteasome for an otherwise UBL-independent FAT10 turnover. The role of Nub1’s UBL domain is thus to localize the Nub1/FAT10 complex to the proteasome for FAT10 engagement and degradation, with no significant allosteric effects, which also agrees with our structural data presented below.

### Nub1’s UBL domain binds to the proteasomal Rpn1 for FAT10 delivery and degradation

To elucidate the structural details of Nub1-mediated FAT10 delivery to the proteasome by cryo-EM, we incubated an excess of pre-formed Nub1/FAT10-Eos complex with ATP-hydrolyzing *hs2*6S proteasome for either 30 s or 60 s before freezing and collecting data. Consistent with actively degrading samples, we observed the proteasome in a non-processing, engagement-competent state with the Nub1/FAT10-Eos complex bound to its surface, and in processing states, where the FAT10-Eos substrate was engaged by the ATPase motor and partially threaded through the central channel (Figure 5A, Figure S12, Table S2). Non-processing and processing conformations of the proteasome could be easily distinguished based on the major conformational transitions that occur upon substrate engagement, wherein the N-ring and AAA+ ATPase ring of the 19S RP become coaxially aligned with the 20S core peptidase ^54–56^. Although we identified several sub-states of the processing proteasome that showed the ATPase hexamer in different nucleotide occupancies and vertical spiral-staircase registers of Rpt subunits, we focus for this study on only a couple of structures that highlight the features important for Nub1-dependent FAT10 degradation.

**Figure 5:**
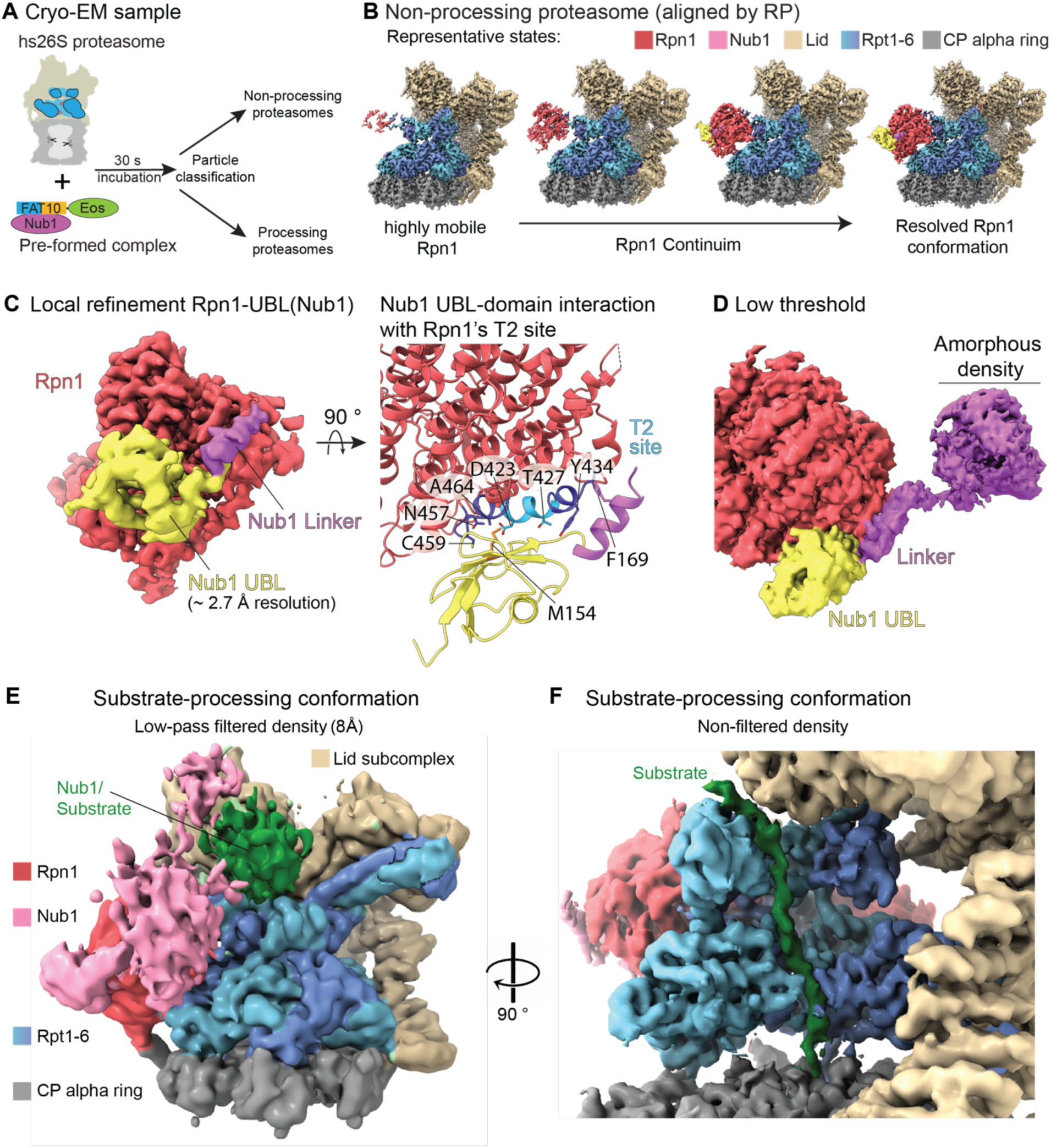
Structures of the Nub1/FAT10-bound *hs*26S proteasome. **A**) For the cryo-EM sample hs26S proteasome was mixed with preformed Nub1/FAT10-Eos complex and incubated for 30 s before freezing. Nub1/FAT10-Eos-bound proteasome particles were then classified into non-processing and processing states. **B)** Representative states for the non-processing proteasome reveal the high flexibility of Rpn1 and consequently the Rpn1-bound Nub1/FAT10-Eos complex. Proteasomes were classified based on Rpn1 conformations, showing a continuum of dynamic and more rigid states. This classification yielded a high-resolution reconstruction for the entire 19S RP, Rpn1 (red), and the Rpn1-bound UBL domain of Nub1 (yellow). **C)** Left: Local refinement of Rpn1 in the non-processing *hs*26S proteasome shows Nub1’s UBL domain (yellow) and the neighboring linker (purple) bound to Rpn1 (red). Right: Local refinement of the Rpn1-Nub1^UBL^ portion of the non-processing *hs*26S proteasome particles allows unambiguous atomic modelling, showing the interactions of a beta sheet in Nub1’s UBL domain (yellow) with the T2 site of Rpn1 (red) and the docking of F169 in Nub1’s UBL linker (purple) with a hydrophobic pocket of Rpn1. **D)** Low-threshold representation reveals an amorphous, poorly resolved density (purple) that likely represents the flexible core body of Nub1 with bound FAT10-Eos. **E)** Low-pass-filtered (8 Å) cryo-EM map of the substrate-processing *hs*26S proteasome that was established by incubating proteasomes with Nub1 and FAT10-Eos for 60 s prior to freezing. Although highly dynamic and poorly resolved, Nub1 (purple) is still bound at this stage of substrate processing. A more distinct yet low-resolution density (green) at the entrance of the ATPase N-ring potentially represents the tough-to-unfold Eos moiety of the FAT10-Eos substrate after the FAT10 portion has been unfolded and translocated into the central channel. **F)** Non-filtered density of the substrate-processing *hs*26S proteasome, rotated relative to the depiction in panel E) by ∼ 90 ° to the left and with the ATPase subunits Rpt4 and Rpt5 removed for a better view of the central channel. Substrate density continues through the AAA+ motor channel and into the 20S core peptidase, indicating that likely the entire FAT10 portion of the FAT10-Eos substrate had been unfolded and translocated at this stage of substrate degradation.

For the non-processing, engagement-competent state of the proteasome, we observed the Nub1/FAT10-Eos complex highly flexibly bound (Figure 5B). These dynamics likely originate from an intrinsic flexibility within the Nub1 complex itself and from binding to Rpn1, which is the most mobile subunit of the 19S RP (Figures S12-S14). After focused 3D classification on Rpn1, we determined 10 structures with resolutions for the ATPase motor ranging from 2.5 – 3 Å and Rpn1 differentially well resolved and at variable positions or angles relative the motor (Figure S13). We chose one representative high-resolution model for this non-processing proteasome that was overall well resolved to ∼ 2.5 Å (Figure 5B, Figure S13) and showed the ATPase hexamer with five ATP and a single ADP, present in Rpt6 (Figure S15). To improve model building for Nub1, we performed a local refinement of the flexible Rpn1, which provided a 2.7 – 3 Å map and allowed us to build an atomic model of Nub1’s UBL domain bound to Rpn1’s T2 site (Figure 5C, S13B). Similar to the deubiquitinase Usp14 (Ubp6 in yeast), whose UBL domain binds to the T2 site of Rpn1 in a slightly varied position ^57^, Nub1’s UBL domain utilizes a hydrophobic center (M154 and L156) that is flanked on either side by charged residues for interactions with Rpn1 (Figure 5C, Figure S16). Interestingly, there is an additional anchor point between Nub1’s F169, located in the linker following the UBL domain, and a pocket on Rpn1 (Figure 5C, Figure S16), which may control the orientation of Nub1 during FAT10 delivery to the proteasomal ATPase pore. However, despite extensive 3D classifications, 3D variability analyses, and local refinements of the 19S RP, we were unable to resolve Nub1’s core, UBA1-3, and the bound FAT10 well enough for fitting individual domains (Figure 5D, Figure S14). Based on the amorphous density observed for the Nub1/FAT10-Eos complex, we assume that these domains get splayed out into various conformations when Nub1 binds to the proteasome or during FAT10 release for proteasomal engagement. High mobility and dynamics of the Nub1 core are likely required for allowing FAT10’s UBL1 domain to sample various positions and orientations during its insertion into the ATPase channel. Ubiquitinated substrates have similarly strong flexibility prior to and during their engagement by the ATPase motor, which so far prevented their visualization by cryo-EM.

For both the 30 s and 60 s time points, we solved actively processing conformations of the Nub1/FAT10-Eos bound *hs*26S proteasome, which showed substrate density in the central channel, reaching through the N-ring, the ATPase ring, and into the degradation chamber of the 20S core peptidase (Figure 5E,F, Figures S17-S19). The length of this substrate trace suggests that in addition to the unfolded UBL1 domain of FAT10, UBL2 was pulled into the central channel. At the entrance to the N-ring, we observed unresolvable yet more defined globular density that may represent the tough-to-unfold Eos domain of the FAT10-Eos fusion at the 60 s time point. Interestingly, we found that Nub1’s UBL domain was still bound to Rpn1’s T2 site at this stage of substrate processing (Figure 5E, S17), indicating that Nub1 is retained at the proteasome even after FAT10 was unfolded and threaded by the ATPase motor. This observation is consistent with a Nub1-mediated delivery and unfolding model by which the entire substrate, including any FAT10-attached protein, must get pulled through the intrinsic loop formed by Nub1’s core and UBA domains. Besides Nub1’s UBL domain stably interacting with Rpn1, there appear to be no other contacts that persist long enough for high-resolution observation by cryo-EM (Figure S17). This is unlike the deubiquitinase Usp14, which, in addition to its UBL domain binding to Rpn1, is further stabilized by interactions between its catalytic ubiquitin specific protease (USP) domain and the ATPase ring to enable substrate deubiquitination ^57–59^.

## Discussion

Here we elucidated a previously unknown mechanism for substrate delivery to the *hs*26S proteasome, in which the cofactor Nub1 acts as an ATP-independent chaperone to trap the UBL1 domain of FAT10 in an unfolded state, recruit it to the 19S RP through binding of the Rpn1 subunit, and present the unstructured N-terminus for engagement by the proteasomal ATPase motor (Figure 6). This cofactor-mediated delivery of a ubiquitin-like modifier for degradation initiation shows fascinating parallels with the mechanisms of the Ufd1/Npl4 cofactor-mediated unfolding of poly-ubiquitinated proteins by the AAA+ unfoldase Cdc48 in yeast (or p97 / VCP in mammals). There, the cofactor subunit Npl4 binds and traps an unfolded ubiquitin moiety of a ubiquitin chain and allows the flexible N-terminus of this initiator ubiquitin to enter the hexameric Cdc48 motor for engagement, subsequent unfolding of the ubiquitin chain, and complete processing of the attached substrate ^6–8^. This mechanism represents a universal mode of delivery that is independent of any substrate features, because ubiquitin provides both the binding sites for Npl4-dependent recruitment and the disordered region for initiation by the Cdc48 unfoldase. Similarly, the UBL1 domain of FAT10 plays both roles in recruitment and initiation, and thus enables the degradation of any FAT10-ylated protein by the 26S proteasome in a Nub1-dependent manner, whereas ubiquitinated substrates of the proteasome require an intrinsic unstructured initiation region of sufficient length and complexity for proteasomal engagement. Since FAT10’s UBL1 domain enters the proteasome first, the location of a FAT10-ylated lysine within a substrate determines from which point the substrate is unfolded and translocated. When reaching the substrate itself, the proteasome will therefore have to translocate a branch point and subsequently two strands in its central channel. However, very little is known about this process, in part because FAT10 conjugation has so far not been successfully reconstituted *in vitro*, and future studies will have to investigate these details of degradation for FAT10-ylated substrates.

**Figure 6:**
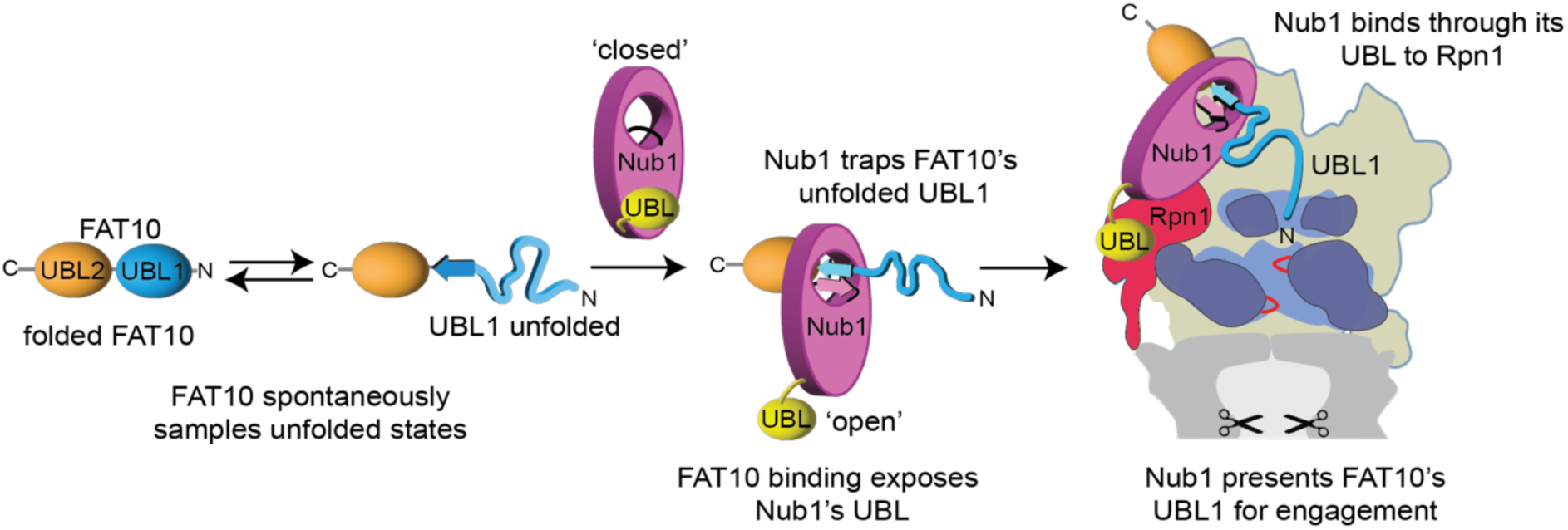
Model for the Nub1-mediated delivery of FAT10-ylated substrates to the 26S proteasome. **A**) FAT10’s low thermodynamic stability allows it to sample partially unfolded states. Nub1 uses a flexible beta-strand linker that lines an internal channel within Nub1 to form a short antiparallel beta sheet with the last beta strand in FAT10’s UBL1 domain and thereby trap the unfolded state. This complex formation is rate-limiting for FAT10 degradation by the proteasome, as it depends on spontaneous FAT10 unfolding. Trapping of FAT10 causes Nub1’s own UBL domain to adopt an open conformation, which enables its interaction with the T2 site of the proteasomal Rpn1 subunit and the presentation of FAT10’s unfolded UBL1 domain for engagement by the ATPase motor.

Isolated FAT10 or FAT10-ylated proteins by themselves are not susceptible to robust proteasomal degradation, and their dependence on the interferon-inducible Nub1 cofactor likely adds an important layer of regulation for fine-tuning the turnover of hundreds of substrates with roles in cell cycle control, NF-κB activation, DNA damage response, autophagy, and mitophagy ^60–64^. Furthermore, there may be other mechanisms to circumvent this dependence on Nub1, for instance through the ubiquitination of FAT10 and its delivery to p97. Interestingly, the Nub1-mediated degradation of FAT10 also has similarities to the recently identified delivery of transcription-factor substrates by the midnolin cofactor for proteasomal turnover. There, the catch domain of midnolin captures a beta strand of the substrate for delivery to the proteasome by a yet unknown mechanism that does not seem to involve midnolin’s UBL domain ^19^.

Our HDX-MS experiments, mutagenesis, biochemical studies, and AlphaFold structural modeling revealed a peculiar mode of complex formation between Nub1 and FAT10, whereby FAT10’s UBL1 domain is inserted into a looped-out portion of Nub1 to form an antiparallel beta sheet that traps this domain in an unfolded state. Essential and apparently rate-determining for this complex formation is the spontaneous unfolding of the UBL1 domain. In the Nub1/FAT10 complex, most of the unfolded UBL1 domain resides on one side of Nub1 for presentation to the proteasomal ATPase motor, whereas the folded UBL2 domain and any conjugated substrate are located on the other side until the proteasome applies mechanical force. The entire substrate may then get threaded through the looped-out portion of Nub1, as suggested by our cryo-EM studies of the actively degrading proteasome that showed Nub1 still bound and potentially interacting with the Eos moiety, after the FAT10 portion of a FAT10-Eos fusion substrate was already translocated. Importantly, Nub1 is highly dynamic and therefore not resolvable, potentially because its domains detach from each other and adopt various different states and positions when bound to the proteasome. Similar to the deubiquitinase Usp14, Nub1 uses an N-terminal UBL domain to interact with the T2 site of the proteasomal subunit Rpn1, but there are no additional persisting contacts to further stabilize the Nub1/FAT10 complex on the proteasome surface. The consequently high mobility seems important for allowing FAT10’s unstructured N-terminus to find and enter the central channel of the ATPase motor.

In summary, we identified the trapping of FAT10’s unfolded UBL1 domain by Nub1 as an elegant principle for substrate delivery and proteasomal engagement. Future studies will have to address whether Nub1 can similarly trap NEDD8 and possibly other beta-strand-containing proteins for processing by the 26S proteasome or p97. The high promiscuity of this Nub1-mediated substrate turnover and FAT10’s expression in immune cells, upon inflammation, viral infection, and in multiple cancers could make the specific FAT10-ylation of neo-substrates for proteasomal degradation an attractive alternative to the proteolysis-targeting chimera (PROTAC) technology, which typically relies on small-molecule induced ubiquitination and frequently requires p97 to prepare well-folded proteins for proteasomal engagement.

## Methods

### Cloning

All truncations were made by using NEBuilder^®^ HiFi DNA assembly master mix (M5520AVIAL, New England Biolabs, NEB) or using Q5 mutagenesis (E0555L, NEB). Point mutations were made using Q5 PCR mutagenesis.

Nub1 sequence was synthesized for codon optimized *E. coli* expression as 3 dsDNA fragments (Integrated DNA technologies, IDT) and assembled into a pGEX-6P-1 vector using NEBuilder^®^ HiFi DNA Assembly Master Mix. The final expressed protein is GST-3C-Nub1, whereby the GST can be removed by precision protease leaving a GPGS overhang at the N-terminus. Nub1L was also cloned the same way as Nub1.

Amino acid sequence for wild-type Nub1:

MAQKKYLQAKLTQFLREDRIQLWKPPYTDENKKVGLALKDLAKQYSDRLECCENEVEKVIEEIRCKAIERGTGNDNYRTTGIATIEVFLPPRLKKDRKNLLETRLHITGRELRSKIAETFGLQENYIKIVINKKQLQLGKTLEEQGVAHNVKAMVLELKQSEEDARKNFQLEEEEQNEAKLKEKQIQRTKRGLEILAKRAAETVVDPEMTPYLDIANQTGRSIRIPPSERKALMLAMGYHEKGRAFLKRKEYGIALPCLLDADKYFCECCRELLDTVDNYAVLQLDIVWCYFRLEQLECLDDAEKKLNLAQKCFKNCYGENHQRLVHIKGNCGKEKVLFLRLYLLQGIRNYHSGNDVEAYEYLNKARQLFKELYIDPSKVDNLLQLGFTAQEARLGLRACDGNVDHAATHITNRREELAQIRKEEKEKKRRRLENIRFLKGMGYSTHAAQQILLSNPQMWWLNDSNPETDNRQESPSQENIDRLVYMGFDALVAEAALRVFRGNVQLAAQTLAHNGGSLPPELPLSPEDSLSPPATSPSDSAGTSSASTDEDMETEAVNEILEDIPEHEEDYLDSTLEDEEIIIAEYLSYVE NRKSATKKN*

Nub1 constructs cloned in this study (truncations and Mutants) are based on Nub1 numbering for the mentioned sequence. The C-terminal potion of Nub1 following on from the NTD-UBL-linker is shortened to trap domain unless referring specifically to UBA domains. Constructs are as follows: UBA1-3 domains, residues 376-528; Nub1 NTD domain, 1-72; UBL domain, 75-161; NTD-UBL, 1-158; Linker-trap domain, 159-601; Linker-trap domain, 175-601; trap-domain, 229-601; ΔUBA1-3, Δ379-527, ΔUBA1-3 + linker, Δ379-527 with insertion at deleted position 4x TGS; ΔUBA1, Δ376-412; ΔUBA2-3, Δ422-527; ΔUBA2-3 + linker, Δ422-527 with insertion at deleted position 4x TGS; ΔBS-linker, 211-223. Mutations in Nub1 constructs: Y572A, D278A, N279A, L562S/I565S, R414A/L418A, D214P/D216P, C317A/R324A. Insoluble Nub1 constructs: Residues 1-372, 228-372, 239-532, 376-601, 376-601. Mutations that gave insoluble constructs: NTD-UBL fragment with M154R and L156R; full length Nub1 with L256D and 260D.

FAT10 was synthesized for codon optimized *E. coli* expression assembled into pCDB179 (Addgene, plasmid number #91960) by Gibson assembly. For labelling with Sortase, a ‘GG’ was added to the N-terminus WT FAT10 sequence by Q5 mutagenesis, whereby after Smt3 cleavage with His-ULP1 a GG scar is left. His-Smt3-Cysless GG-FAT10 (FAT10^C0^) was gene synthesized and subcloned into a pET28 vector by GeneArt (Thermo Fisher Scientific), the mutations are C7T, C9T, C134L, C160S and C162S. GG-FAT10 was used for comparison to GG-FAT10^C0^, GG-FAT10^H75A^, GG-FAT10^H75D^, GG-FAT10^H75K^ in assays. For all other experiments involving unlabeled FAT10, WT FAT10 sequence displayed below was used.

Amino acid sequence for wild-type FAT10:

MAPNASCLCVHVRSEEWDLMTFDANPYDSVKKIKEHVRSKTKVPVQDQVLLLGSKILKPRRSLSSYGIDKEKTIHLTLKVVKPSDEELPLFLVESGDEAKRHLLQVRRSSSVAQVKAMIETKTGIIPETQIVTCNGKRLEDGKMMADYGIRKGNLLFLACYCIGG*

For creating Eos3.2 constructs, Ub_4_-Eos3.2-intein-CBD (chitin-binding domain) and Ub_4_-Eos3.2-tail-intein-CBD from a previous publication ^5^ were used as templates. For FAT10-Eos, FAT10-Eos-tail, FAT10^ΔUBL2^-Eos, FAT10^ΔUBL2^-Eos-tail, FAT10^ΔUBL1^-Eos, and FAT10^ΔUBL1^-Eos-tail, the wild type FAT10 vector (His-Smt3-FAT10) was linearized, and PCR fragments of Eos-intein-CBD or Eos-tail-intein-CBD were inserted via HiFi assembly. Using the FAT10-Eos and FAT10-Eos-tail vectors as templates, they were linearized by PCR, removing FAT10 and replacing with Nub1’s UBL domain (which were amplified from full length codon optimized Nub1 sequence) or NEDD8 (ordered as a dsDNA from IDT, codon optimized for *E.coli*) by HiFi assembly.

The plasmid for expression of His-ULP1 was from Addgene (Plasmid #64697), the His-Sortase plasmid was created as previously described ^4^, and the His-GST-3C plasmid as well as purified His-TEV protease were sourced from QB3 MacroLab (UC Berkeley).

### Protein expression in E. coli

All proteins were expressed in BL21* *E. coli* cells grown at 37°C with 200 rpm in TB medium (24g yeast extract, 20g tryptone, 8 mL glycerol, and buffer with phosphate pH7.2). After letting cells cool down to ∼16°C, expression was induced after reaching OD_600nm_ 0.6-0.8 with IPTG (0.25 µM), and cells were left growing overnight at 16°C.

### Nub1 purification

After overnight expression at 16°C, all Nub1 expressing cells are harvested and suspended in lysis buffer (60 mM HEPES pH 7.4, 25 mM NaCl, 25 mM KCl, 5% (v/v) glycerol, 10 mM MgCl_2_, 0.5 mM TCEP) supplemented with EDTA-free protease inhibitor tablets (11836170001, Roche) and benzonase (70664, Novagen®). Cells are lysed by sonication (on ice) and clarified at 15,000 x g for 45 min at 4°C. Lysates are then flowed slowly (∼1 mL/min) by gravity over pre-equilibrated (in lysis buffer) GSH resin (16101, ThermoFisher Scientific) multiple times, before successive washes (at least 10 column volumes (cv) of lysis buffer) and suspension in 2 cv of lysis buffer. GST-3C protease was added for overnight incubation at 4°C before collection of flow-through (followed by collection of another two cv washes over resin), clarification (4,000 x g, 15 min) and concentration using an Amicon Ultra-15 30-kDa cut off concentrator (fUFC905008, MilliporeSigma) or gel filtration by a Superdex (SD)200 increase 10/300 column or SD200 16/600 column, depending on scale and yields of protein. Fractions containing Nub1 were collected from a single peak, concentrated to ∼10-15 mg/mL and snap-frozen as single use aliquots (10 µL) in liquid N_2_ for storage at –80°C. Protein concentration was estimated using A_280nm_ and all Nub1 proteins had a A_280/260nm_ ratios between 0.5-0.6.

### FAT10 purification

His-Smt3-FAT10 expressing cells were harvested and suspended lysis buffer supplemented with benzonase, EDTA-free protease inhibitor tablets, 300 mM NaCl and 20 mM imidazole. Cells were sonicated and clarified before flowing lysate over pre-equilibrated Ni-NTA resin several times. Contaminants were removed by several successive washes with lysis buffer, before incubation of resin with His-ULP1 protease overnight in lysis buffer supplemented with 150 mM NaCl, the cleavage reaction was not mechanically moved and instead resuspended with a pipette a few times before leaving the reaction overnight at 4°C. The reaction was typically mixed one more time before moving to room temperature for 10 mins. After this, flowthrough containing cleaved FAT10 was collected for anion exchange. FAT10 was carefully diluted with lysis buffer (usually about 10-fold in volume) before binding to a HiTrap SP HP column (17115201, Cytiva) and elution over a linear gradient (0-1000 mM NaCl). FAT10 eluted as a single peak, which was concentrated using an Amicon Ultra-15 10-kDa cut off concentrator, clarified by centrifugation (20,000 x g at 4°C) before fractionation with an SD75 increase 10/300 column or SD75 16/600 column. FAT10 was eluted as a single peak and was concentrated to ∼10 mg/mL, before flash freezing as single use 10 µL aliquots and storage at –80°C. All FAT10 mutans were purified as per wildtype, except for the single UBL1 domain of FAT10, which skipped the cation exchange, and His-Smt3-FAT10 fusions, which were eluted with lysis buffer supplemented with 250 mM imidazole.

### Purification of UBL-Eos3.2

All UBL-Eos3.2 constructs were expressed as fusion proteins with an N-terminal His-Smt3 and C-terminal intein-CBD. Cells were lysed in lysis buffer supplemented with benzonase, EDTA-free protease inhibitor tablets and 20 mM imidazole before sonication, clarification, and binding over Ni-NTA resin. After extensive washing with lysis buffer, proteins were eluted with lysis buffer + 250 mM imidazole and bound to chitin-resin (S6651L, NEB), before washing in lysis buffer and overnight incubation with lysis buffer + 200 mM DTT and his-ULP1 protease. The flowthrough was collected and flowed by gravity over Ni-NTA resin to remove His-ULP1 and His-Smt3 proteins before concentration and gel filtration using an SD200 16/600 column in GF buffer. Protein concentrations were estimated by absorbance at A507 nm and A280 nm, since not all Eos3.2 matures, we used the concentration estimated from A280 as the concentration of UBL-Eos3.2 substrates.

### Purification of Sortase, His-ULP1, and His-GST-3C

His-SortaseA and His-ULP1 were purified with identical conditions, after suspension in lysis buffer supplemented with benzonase, 200 mM NaCl and 20 mM imidazole, cells were lysed by sonication and clarified, before flowed over Ni-NTA resin by gravity. Ni-NTA resin was washes extensively and proteins were eluted with lysis buffer supplemented with 250 mM Imidazole and 150 mM NaCl. For His-GST-3C protease, conditions were similar except lysate was flowed over GSH-resin before washing and elution with 20 mM GSH in lysis buffer. Eluted His-GST-3C was subsequently bound to a HiTrap Q HP column (17115401, Cytiva) and eluted over a linear gradient (0-1000 mM NaCl). All proteins were concentrated and fractionated using an SD200 16/600 column in 30 mM HEPES pH7.4, 150 mM NaCl, 5% (v/v) glycerol and 0.5 mM TCEP. Proteins were concentrated to ∼10 mg/mL (estimated by A280nm) and frozen in liquid N_2_ for storage at – 80°C.

### Human 26S proteasome purification from HTBH-Rpn11 expressing HEK293 cells

The hexahistidine, TEV cleavage site, biotin and hexahistidine (HTBH)-tagged human 26S proteasomes expressing HEK293 cells were previously generated ^65^ and a kind gift from L. Huang. Cells were adapted for suspension to increase scale and ultimately yields of *hs*26S proteasome. For adaptation, cells were grown by gradually lowering Fetal Bovine Serium (FBS, 16000044, ThermoFisher Scientific) concentration from 10%-5% (v/v) on plates, and after 3 passages were grown in FreeStyle™ 293 Expression Medium (12338018, ThermoFisher Scientific) with 2% (v/v) FBS. Cells were harvested and moved to a shaker flask, where the suspension cells were grown at 8% CO_2_ 37°C with 120 rpm shaking in FreeStyle™ 293 Expression Medium with 2% (v/v) FBS. Cells were passaged twice a week at 5 × 10^5^ and newly thawed cells were grown with puromycin.

For purification, 4L of HTBH-Rpn11 HEK293 cells were grown for 72 hours after passaging to 5 × 10^5^ and harvested by centrifuge at 4,000 x g. Cell pellets were resuspended in lysis buffer supplemented with benzonase, EDTA-free proteasome inhibitor tablets, 0.01% NP-40 and 2 mM ATP. Cells were lysed by a Dounce homogenizer (usually 15X) followed by sonication on ice with low amp (20%) and 10 seconds on and off for 2 min. Lysates were clarified for 60 min at 4°C and flowed over pre-equilibrated Pierce™ High-Capacity Streptavidin Agarose (3 mL of resin). After 2X rounds of binding, beads were carefully sequentially washed with lysis buffer (+2 mM ATP) 5X (3 mL) before suspension in 3 mL of lysis buffer (+2 mM ATP). TEV protease (250 µg) was added, and resin was incubated at room temperature for 60 min or overnight at 4°C. Flowthrough from resin is collected, beads were washed with 2 additional column volumes and collected for concentration to ∼250 µL with an Amicon Ultra-15 100-kDa cut off concentrator. The sample was clarified at 20,000 x g for 15 min at 4°C before fractionation with an S6 increase 10/300 column in GF buffer supplemented with 2 mM ATP. Fractions containing 26S proteasomes were pooled and concentrated to ∼50-100 µL before freezing as 10 µL single use aliquots in liquid N_2_ and stored at –80°C. Concentration of samples was measured by Bradford reagent with BSA as a standard, we assumed that the *hs*26S proteasome is 2600 KDa for molar calculations.

### Purification of sc26S proteasome and sc20S proteasome

The *S.cerevisiae* 20S core and 26S holoenzyme was purified from the yAM54 (MATa his3Δ200, leu2-3,112 lys2-801 trp1Δ63 ura3-52 PRE1-3xFLAG::Ylplac211(URA3) or YYS40 (MATa ade2-1 his3-11,15 leu2-3,112 trp1-1 ura3-1 can1-100 RPN11:RPN11-3XFLAG (HIS3)) yeast strains, respectively. Yeast were grown in YPD (yeast extract, peptone and dextrose) at 30°C for 3 days before harvesting. Briefly, yeast were lysed by freezing in lysis buffer with liquid nitrogen and using a 6875 Freezer Mill Dual Chamber Cryogenic grinder (SPEX Sample Prep). The 3xFLAG-Pre1 yeast cells were resuspended in 60 mM HEPES, pH7.6, 500 mM NaCl, 0.1 % (v/v) NP40, 5% glycerol (v/v) and for 3xFLAG-RPN11 are lysed in 60 mM HEPES, pH7.4, 25 mM NaCl, 25 mM KCl, 5% (v/v) glycerol, 10 mM MgCl_2_ 0.5 mM TCEP and 2 mM ATP. Proteasomes were bound to M2anti FLAG resin (Sigma), where 20S particles were washed with 1 M NaCl to remove bound regulatory particle, and 26S is washed only in low salt with 60 mM HEPES, pH7.4, 25 mM NaCl, 25 mM KCl, 5% (v/v) glycerol, 10 mM MgCl_2_ 0.5 mM TCEP and 2 mM ATP. Both complexes are eluted from FLAG-resin using 0.3 mg/mL 3 x FLAG peptide, and further purified by size-exclusion chromatography with a Superose6 Increase 10/300 column (GE) equilibrated 60 mM HEPES, pH7.4, 25 mM NaCl, 25 mM KCl, 10 mM MgCl_2_ 2, 5% (v/v) glycerol and 0.5 mM TCEP (for 26S 2 mM ATP was also added). Proteins are quantified using Bradford reagent with BSA as a protein standard.

### Degradation assays by SDS-PAGE

All SDS-PAGE degradation assays were performed at 30°C in 0.65 mL microcentrifuge tubes (1605-0001, SealRite). Proteins were diluted with reaction buffer (60 mM HEPES pH 7.4, 25 mM NaCl, 25 mM KCl, 5% (v/v) glycerol, 10 mM MgCl_2_, 0.5 mM TCEP and 0.5 mg/mL BSA) supplemented with 5 mM ATP and 1X ATP regeneration mix (5 mM ATP, 0.03 mg/ml creatine kinase and 16 mM creatine phosphate). FAT10 alone or FAT10 with excess Nub1 are incubated in reaction buffer (in a 2 to 4x final assay concentration) for at least 30 mins on ice before diluting with buffer and mixing with 2x 26S proteasome stock. Concentrations used in specific assays are indicated in figure legends but for example in Figure 1B, Nub1 at 30 µM was incubated with FAT10 at 10 µM for 30 min on ice before mixing with 2X 26S proteasome (2X is at 200 nM) to reach a final rection volume of 10 µL with 100 nM 26S proteasome, 15 µM Nub1 and 5 µM FAT10. After incubation for an indicated amount of time, reactions were quenched with 10µL SDS-PAGE loading buffer (Tris-base to pH 7, 20% (v/v) glycerol, 4% (w/v) SDS and 0.04 mg brilliant blue, 200 mM DTT) and 12 µL was loaded onto SDS-PAGE gels for Coomassie staining.

### Multi-turnover FAT10-Eos3.2 degradation assays

For multi-turnover reactions, FAT10-Eos3.2 were always at higher concentrations than 26S proteasome, specific concentrations of each component are in Figure legends. Degradation of Eos3.2 was monitored by SDS-PAGE or loss in fluorescence loss using a BMG Labtech CLARIOstar plate reader at 30°C with emission 520 nm after excitation with 500 nm. Proteins were diluted in pre-warmed reaction buffer with 5 mM ATP and preformed FAT10-Eos3.2+Nub1 complexes were mixed 1:1 with 26S proteasomes to reach final concentrations. For Michaelis-Menton kinetics, the initial rate at each FAT10-Eos3.2 concentration was fit and converted to degradation rate (substrate/Enzyme/min^−1^) before plotting and calculating K_Cat_ and K_m_ using GraphPad (Prism). Individual replicates are processed and the mean +/− SD of the K_Cat_ and K_m_ is reported.

### Single turnover kinetics

For single turnover experiments the unfolding rate of Eos3.2 was measured by incubation with excess hs26S proteasome and Nub1. For Figure 1G and 1H, fluorescence loss of Eos3.2 was measured using a BMG Labtech CLARIOstar plate reader at 30°C with emission 520 nm after excitation with 500 nm. Samples were first loaded at 2X concentrations in separate wells of a pre-warmed 384-well plate. After 1 min of incubation in the plate reader, 5 µL of hs26S proteasome (4 µM) was pipetted from one well to the substrate containing well and mixed quickly before starting the read, in this way the first 6-8 seconds was usually not recorded as it takes time for the machine to begin reading. Typically, reactions were monitored for ∼20 min before fitting single or double exponentials using GraphPad (Prism). The single exponential K or double exponential K_fast_ was used to calculate the unfolding rate (substrate/enzyme/min^−1^) and the mean +/− SD of three experiments if reported for each condition.

### Sortase labelling

His-Sortase at 25 µM was incubated with FAT10 (with a GG at its N-terminus) at 30 µM for 30 mins at room temperature with 5-FAM conjugated to the N-terminus of HHHHHHLPETGGG peptide (ordered from Biomatik) at 500 µM in 1X reaction buffer without BSA, supplemented with 10 mM CaCl_2_ and 1 mM DTT. Labeled FAT10 (^FAM^FAT10) proteins were enriched using Ni-NTA resin followed by spin filtering through 0.22 µM spin columns and fractionation using a SD75 increase 10/300 column. ^FAM^FAT10 concentrations are estimated using FAM absorbance (5-FAM=A492nm).

### Binding assays with ^FAM^FAT10

All binding assays were performed in 10 μL volumes in a preheated (30°C) 384-well black plate (Costar) using a BMG Labtech CLARIOstar plate reader. Polarization of ^FAM^FAT10 was measured by with excitation at 480 nm and emission at 535 nm. For measuring the K_D_ between FAT10 and Nub1, ^FAM^FAT10 was incubated at 10x final concentration, either alone, or with increasing Nub1 concentrations for 30 mins on ice, before diluting to 1X (100 nM ^FAM^FAT10 + Nub1 at indicated concentrations) and measuring polarization, the end point with stable signal was taken as the final mP value. For measuring kinetics of binding (K_on_), ^FAM^FAT10 at 2x concentration (40 nM) was mixed with Nub1 at 2x concentration and polarization was measured over time. For single concentrations, binding experiments, at 2x ^FAM^FAT10 (200 nM) were pre-incubated with 2x Nub1 (10 µM) for 30 mins on ice before measuring polarization and taking the end point after polarization stable. GraphPad (prism) was used for data analysis. For K_on_ single exponentials were fit to each curve and K was taken to fit a linear regression with the gradient as K_on_, data shows mean +/− SD (n=3).

### ATPase assays

ATP hydrolysis rates were determined using NADH depletion, where absorbance at 340 nm was monitored over time in a BMG Labtech CLARIOstar plate reader at 30°C. The 1x ATPase mix 5 mM ATP, 3 U mL^−1^ pyruvate kinase (Sigma), 3 U mL^−1^ lactate dehydrogenase (Sigma), 1 mM NADH, 7.5 mM phosphonyl pyruvate (Sigma) was incubated with hs26S proteasome (typically at 100 nM) in the presence or absence of FAT10 (10 µM) and or Nub1 (15 µM) in reaction buffer. When using FAT10+Nub1, proteins were incubated on ice for at 20 mins. ATPase rates are measured using the linear part of the curve.

### Peptidase assays

Stocks of each reagent were prepared at 4x final concentrations: LLVY-AMC (400 µM), *hs*26S proteasome stock (400 nM), FAT10 (60 µM), Nub1 (60 µM) in reaction buffer supplemented with 5 mM ATP. For *hs*26S proteasome stocks, 4X ATP regeneration mix was also supplemented, and in conditions with ATPγS at 20 mM was added to the 4X *hs*26S stock. FAT10 and Nub1 were diluted 1:1 with buffer or each other to form a complex on ice for 30 mins, making a 2X substrate stock. Samples (pre-warmed to 37°C) were mixed in the following order, FAT10 (or Nub1, or FAT10+Nub1) followed by LLVY-AMC, followed by *hs*26S proteasome to initiate the reaction. AMC-fluorescence was measured at 445 nm after excitation at 345 nm in a preheated 384-well black plate (Costar) using a BMG Labtech CLARIOstar plate reader. The linear part of the reaction was extracted and converted to a percentage relative to the 26S proteasome alone.

### Size exclusion chromatography (SEC) and SEC-Multi angle light scattering (MALS)

Nub1 (50 µM) was incubated with FAT10 (75 µM) for 30 mins on ice before fractionating with an SD200 Increase 10/300 column. A single peak for the Nub1/FAT10 complex was pooled and SEC-MALS were conducted on Agilent Technologies 1100 series with a 1260 Infinity lamp, Dawn Heleos II and the Optilab T-Rex (Wyatt Technologies), with an SD200 Increase 10/300 column. The column was equilibrated with 60 mM HEPES pH7.4, 50 mM NaCl, 50 mM KCl, 5% glycerol, 10 mM MgCl_2_ and 0.5 mM TCEP with a flow rate of 0.5 mL/min. The same was repeated for Nub1 alone.

### HDX-MS sample preparation

Samples for FAT10 and Nub1 were diluted to 10X stock concentration, so that 4 stocks of protein were prepared: FAT10 (10 µM), FAT10+Nub1 (10 µM + 15 µM), Nub1 (10 µM) and Nub1+FAT10 (10 µM + 15 µM) and samples were left on ice for at least 30 mins. Replicates for each set of HDX-MS experiments were done with three different preparations of FAT10 and Nub1. Labelling buffer was prepared as a 10X stock by diluting 300 mM HEPES pH_read_ 7.0 (effectively pD7.4), 250 mM NaCl, 250 mM KCl, 100 mM MgCl_2_, 5 mM TCEP in 100% D_2_O; an equivalent H_2_O (10X stock) buffer was made at pH7.4. A 2X quench buffer was prepared with 200 mM glycine pH2.4, 3.5 M Guanidium hydrochloride and 200 mM TCEP. Each sample was prepared by diluting 2 µL in 18 µL of D_2_O labelling buffer and incubated for an indicated amount of time at 25°C before quenching (rapidly mixing 20 µL, so 1:1 with quench buffer) with rapid mixing) and flash freezing in liquid N_2_ and storage at –80°C. An unlabeled sample was prepared as above except diluted into H_2_O buffer before quenching.

### HDX-MS

Samples (in a random order) were immediately thawed and injected one-by-one into a cooled valve system (Trajan LEAP) coupled to an LC system (Thermo UltiMate 3000) maintained at 2°C. Proteins were digested on-column by flowing quenched samples at 200 μL/min in 0.1 % formic acid over in-house prepared protease columns (2mm ID × 2 cm, IDEX C-130B) at 10°C. The proteases, aspergillo pepsin (Sigma-Aldrich, P2143) and porcine pepsin (Sigma-Aldrich, P6887), were crosslinked to POROS Al aldehyde activated resin (Thermo Scientific, 1602906) in that order, respectively. Peptides were desalted for 4 minutes with Thermo Scientific POROS R2 reversed-phase resin (Thermo Scientific POROS R2 reversed-phase resin 1112906) hand packed into a trap column (1 mm ID × 2 cm, IDEX C-128) at 2°C. Subsequently, peptides were separated using a C8 column (Thermo Scientific, BioBasic-8 5 μm particle size 0.5 mm ID × 50 mm 72205– 050565) at a flow rate of 40 μL over 14 minutes with a 5-40% gradient of 100% Acetonitrile and 0.1% formic acid followed by 90% over 30 seconds. Peptides were eluted directly into a Q Exactive Orbitrap Mass Spectrometer operating in positive mode (resolution 70000, AGC target 3e6, maximum IT 50 ms, scan range 300–1500 m/z). Prior to all subsequent injections, protease columns were washed 2x with 100 μL 1.6 M Guanidium hydrochloride and 0.1% formic acid. All of the LC and MS methods were performed using Xcalibur 4.1 control software (Thermo Scientific). Analytical and trap column were subject to saw-tooth washes and equilibrated at at 5% of 100% Acetonitrile and 0.1% formic acid. For undeuterated samples and each condition (FAT10, FAT10+Nub1, Nub1 and Nub1+FAT10) a separate MS/MS experiment was run to identify peptide lists using the MS settings described except the following settings: resolution 17500, AGC target 2e5, maximum IT 100 ms, loop count 10, isolation window 2.0 m/z, NCE 28, charge state 1 and ≥7 excluded, dynamic exclusion of 15 seconds.

Byonic (Protein Metrics) was used to identify FAT10 and Nub1 peptides from MS/MS spectra. Peptide lists (sequence, charge state, and retention time) were exported from Byonic and imported into HDExaminer 3 (Sierra Analytics). When multiple peptide lists were obtained, all were imported and combined in HDExaminer 3. HDX-MS peptides were analyzed using HDExaminer 3 where peptide isotope distributions and deuteration amounts are calculated and extracted. For Nub1 all peptides were analyzed using unimodal analysis, whereas a large majority of FAT10 peptides mass spectrum were fit with bimodal which calculates two centroid peaks and therefore two deuterated levels, we did not consider intensities of each peak due to mixed EX1/EX2 deuterated uptake kinetic regimes, making accurate fitting ambiguous when peaks overlapped. We looked for the presence of absence of bimodal distributions and described an overarching effect from Nub1 binding. Uptake plots are fit from experimentally calculated deuterated levels for each peptide at each time point and wood’s plots displaying all peptides were generated by extracting data from HDExaminer and plotting with Jupiter notebook using python and Matplotlib. For FAT10 we generated the wood’s plot by only considering the left peak (lightest peak) with comparison to unimodal peaks or when present the left peaks for bimodal obtained from FAT10+excessNub1 data.

### Cryo-EM Sample preparation and data collection

Samples were diluted in 20 mM HEPES pH7.4, 25 mM NaCl, 25 mM KCl, 5 mM MgCl_2_, 2 mM ATP, 2.5% glycerol and 0.02% NP-40 as 2X stocks and centrifuged at 15,000 x g for 15 min at 4°C. Proteasomes (4 µM) were mixed 1:1 with preincubated (20 min) FAT10-Eos+Nub1 (10 µM + 12 µM) complexes (final 1 µM, 5 µM and 6 µM, respectively) for 30 seconds and 60 seconds before cryo plunging. Samples (3.5 µL) were applied to glow discharged (25 mA, 25 seconds) UltrAufoil^®^ R 2/2, 200 Mesh, Au grids (Q250AR2A, Electron Microscopy Sciences). Using a Vitrobot (ThermoFisher), glow discharged grids were placed at 100% humidity and samples were applied and immediately blotted for 2.5 seconds (10 blot force) before plunge freezing in liquid ethane.

Grids were clipped and transferred to Titan Krios transmission electron microscope operated at 300 KeV (ThermoFisher) with an energy filter (GIF quantum) and equipped with Gatan K3 using serial EM. Images were taken at a nominal magnification of x81,000 (1.048 Å pixel size) in super resolution mode with a defocus ranging from –0.5 – 1.7 µm, using SerielEM ^66^. We collected 50 frames per shot with a total electron dose ∼50 e^-^ Å^-2^s^−1^. A total of 20,565 movies were collected for the 30 second data set and 19,128 movies for the 60 second data set.

### Cryo-EM data processing

All micrographs were patch motion corrected with CTF estimation using CryoSparc v4.3.1 ^67^. From the 30 second data set and 60 second data, 20,565 and 19,128 corrected micrographs were subjected to blob picker. Picked particle blobs were extracted with a 720-pixel box binned by 2. Particles were subjected to multiple rounds of 2D classification before taking a small subset of particles (∼50k) and generating 4 *ab-initio* reconstructions, where a single 30S model with secondary structure features was selected. The 30S *ab-initio* model was used to seed multiple rounds of heterogenous refinement (with 4-10 classes) with binned pixel size of 128. Particles that reached Nyquist frequencies (when binned) in heterogenous refinement were aligned by homogenous refinement in C2 to a symmetry aligned low pass filtered 30S model, which was aligned manually in UCSF Chimera ^68^. Symmetry expansion was used to effectively double the number of particles, as we wanted to focus on features of the 19S. Particles were then shifted using volume alignment tools in CryoSparc to where the 19S was at the center of the box and particles were re-extracted with a box size of 280 pixels. Homogenous reconstruction was used to generate a 19S model with just over half of the 20S, followed by homogenous refinement. The 19S model was used to seed multiple rounds of heterogenous refinement, which results in some low-resolution classes, a few non-processing classes and a single class containing substrate engaged proteasomes. For selected non-engaged and engaged proteasome particle stacks, 20S signal was removed by generating a mask and using particle subtraction. Subtracted particles were subject to homogenous reconstruction and homogenous refinement.

For substrate engaged proteasomes, rounds of heterogenous refinement were used again to separate out states, resulting in 4 major ATPase states, each of which was subjected to homogenous refinement followed by NU refinement ^69^. The largest class of substrate engaged proteasomes from the 30 seconds data set was subjected to another round of 19S masked 3D classification, and NU refinement, resulting in one high resolution representative state for FAT10-engaged Nub1-bound *hs*26S proteasome, and subsequent model building. The above method was also used for the 60 second data set, except the final classification step was omitted. Many attempts were made to resolve additional density found in models including 3D variability analysis ^70^, 3D classification, heterogenous refinement, and particle subtraction.

For particles in non-engaged proteasome stacks, Rpn1 was clearly flexible with extra density. We generated 10 classes through Rpn1 masked 3D classification and performed homogenous refinement on each class. This gave rise to 10 non-engaged proteasome structures with Rpn1 is varied positions, the rest of the 19S models appeared almost identical. Two structures with a total of 103K particles showed Rpn1 as completely mobile, where extra density could still be seen through low pass filtering models. However, we could not resolve additional structures likely due to the continuous mobility of Rpn1. The other 8 classes showed a well resolved Rpn1 with a UBL domain attached at variable resolutions. The highest resolution model was chosen for NU refinement and this mode was used for model building. In addition, the 8 models were combined for local refinement for a single high-resolution model of Nub1’s domain bound to Rpn1, which also allowed further modeling of the linker from Nub1. Using the combined stack, homogenous refinement was again used to align particles but with a larger mask covering more extra density, followed by 3D classification (20 classes with filtering resolution to 15 Å) and homogenous refinement of each class. Each class contained an amorphous mass attached to the UBL of Nub1, but despite effort could not be resolved. We used one of these models to represent the model for how Nub1 is dynamically moving relative to the 19S and its own UBL, likely sampling variable positions to help FAT10 engage the proteasome.

### Cryo-EM Model building and visualization of structures

For non-engaged proteasome models, we used 7W37 ^57^ as a starting model with rigid body fitting using Chimera. However, our high-resolution models allowed us to detect multiple register errors. We replaced several chains using AlphaFold models ^71^, the replaced chains are: A, B, C, D, F, U, V, W, X, Y, Z, a, b, c, d, f and g. We did not replace chain E, G, H, I, J, K, L, M and e. We deleted parts or most of chains N, O, P, Q, R, S, T. Coot was used to manually curate side chain positions and secondary structure differences from AlphaFold models ^72^. The high-resolution data allowed us to build unmodelled sequences, such as the N-terminus of Rpn1, which is sandwiched between the toroidal domain of Rpn1 and Rpt1. We were able to unambiguously nucleotide densities. For engaged proteasome models we used 6MSK (Zhang et al., 2022) with rigid body fitting and extensive remodeling with coot. Real space refinement in Phenix was performed iteratively with model building in Coot ^73,74^. Figures were generated using PyMOL (The Molecular Graphics System, Version 1.8, Schrödinger, LLC; http://www.pymol.org/), UCSF chimera and ChimeraX ^75^. Local resolution was displayed shown for each structure using local resolution estimation in CryoSparc with 0.143 from FSC curves and Chimera with color surface. Low pass filtered models were generated in CryoSparc with volume tools.

## RESOURCE AVAILABILITY

### Lead contact

Further information and requests for resources and reagents should be directed to and will be fulfilled by the lead contact, Andreas Martin.

### Materials availability

All constructs generated in this study are available from the lead contact upon request and completion of a Material Transfer Agreement.

### Data and Code Availability

- All data generated or analyzed during this study are included in this manuscript and the Supplementary materials. Structural data are available in the Electron Microscopy Databank and the RCSB Protein Databank (EMDB ID 42506 and PDB ID 8USB for the non-processing 26S proteasome, EMDB ID 42507 and PDB ID 8USC for the substrate-processing 26S proteasome at 30 s after substrate addition, and EMDB ID 42508 and PDB ID 8USD for focused refinement of the proteasomal Rpn1 subunit with bound UBL domain of Nub1).
- This paper does not report original code.
- Any additional information required to reanalyze the data reported in this paper is available from the lead contact upon request.

## Acknowledgments

We thank all members of the Martin lab for discussion and support. We thank Lan Huang (UC Irvine) for gifting httb-HEK293 cells and the UCB Cell Culture Facility (RRID: SCR_017924) for maintaining the httb-HEK293 cell culture. Cryo-EM data were collected at the UCB Cal-Cryo facility, and we thank Dan Toso and Ravindra Thakkar for cryo-EM operational support.

## Author contributions

CA., K.C.D. and A.M. conceived the study, C.A. and A.M. designed experiments, C.A. cloned constructs, expressed, and purified proteins, and C.A. and K.C.D. performed biochemical measurements as well as data analyses. C.A. performed and analyzed HDX-MS experiments with guidance from S.M.C. and S.M.. S.M.C. prepared pepsin-columns and maintained HDX-MS instrument. C.A. performed cryo-EM sample preparation, data collection, and data processing. C.L.G. and C.A. generated atomic models based on cryo-EM maps. C.A. and A.M. wrote the manuscript with comments from all authors.

## Supplementary Figures

**Figure S1:**
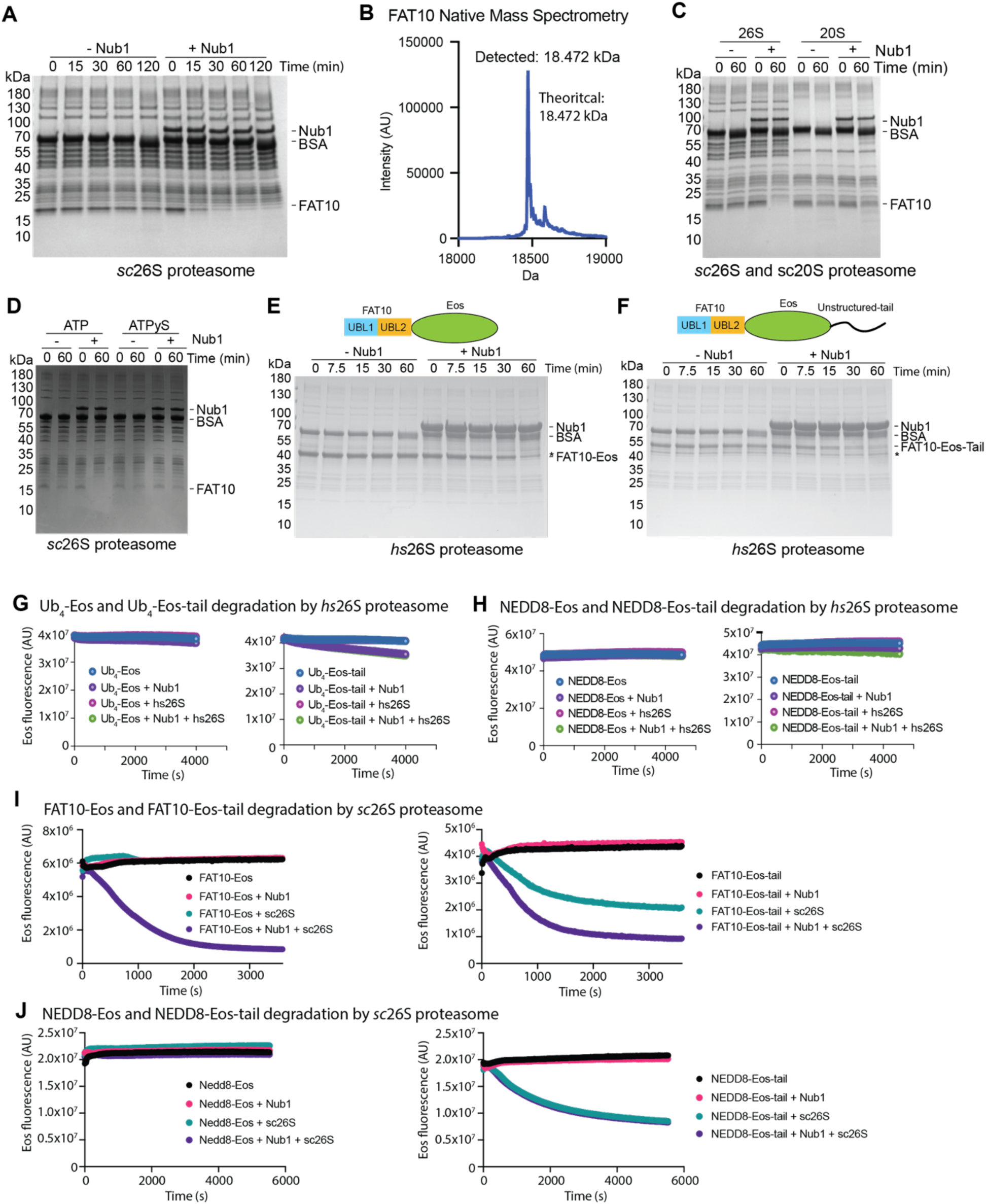
Supporting data for Figure 1; Nub1 accelerates degradation of FAT10 for yeast proteasomes, does not accelerate degradation of Nedd8 fusions, and a disordered tail can bypass the requirement of Nub1 for degradation by yeast proteasomes but not human proteasomes. **A**) Coomassie-stained SDS-PAGE gel showing the time course for the degradation of FAT10 (10 μM) by *sc*26S proteasome (100 nM) with and without Nub1 (5 μM). **B)** Native mass spectrometry confirms the lack of any truncations or modifications for *E. coli*-expressed human FAT10. **C)** Coomassie-stained SDS-PAGE gel to analyze the end points of FAT10 degradation (10 μM) by *sc*26S proteasome (100 nM) or *sc*20S CP (100 nM) in the absence or presence of Nub1 (5 μM). There is only very slow degradation observed for *sc*20S CP, indicating that ATP-hydrolysis driven unfolding and translocation by the 19S RP is required for robust FAT10 degradation. **D)** Same assay as in C) in the presence of either ATP or the slowly hydrolyzed ATP analog ATPγS. The degradation in the presence of ATPγS confirms the dependence of FAT10 processing on ATP-dependent mechanical unfolding and translocation. **E)** Degradation of FAT10-Eos (5 μM) and **F)** FAT10-Eos-tail (5 μM) by *hs*26S proteasome (100 nM) with and without Nub1 (15 μM) demonstrates that FAT10 together with Nub1 can mediated the turnover of tough-to-unfold substrates, but adding an disordered engagement sequence does not bypass the requirement for Nub1. **G)** Example traces showing the loss of Eos3.2 fluorescence during the degradation of Ub_4_-Eos (5 μM) and Ub_4_-Eos-tail (5 μM) by *hs*26S proteasome (100 nM) in the presence or absence of Nub1 (15 μM). While Ub_4_-Eos3.2-tail is robustly degraded with or without Nub1, Ub_4_-Eos is not. **H)** Same fluorescence-based assays as in G) except using NEDD8-Eos and NEDD8-Eos-tail. Neither of those substrate are degraded by *hs*26S proteasome, even in the presence of Nub1. **I)** Left: Degradation of FAT10-Eos (2 μM) by *sc*26S proteasome (100 nM) in the absence or presence of Nub1 (5 μM), showing the Nub1 dependence of FAT10 turnover even for the yeast proteasome. Right: Degradation of FAT10-Eos-tail (2 μM) by *sc*26S proteasomes (100 nM) in the absence or presence of Nub1 (5 μM), indicating that a flexible tail on the substrate can bypass the requirement of Nub1. FAT10 is sufficient for substrate recruitment to *sc*26S proteasomes, while the flexible tail allows engagement by the ATPase motor, which is in contrast to *hs*26S proteasome. **J)** Same fluorescence-based assays as in I), except using NEDD8-Eos and NEDD8-Eos-tail. While NEDD8-Eos cannot be degraded by *sc*26S proteasome, NEDD8-Eos-tail is robustly turned over in a Nub1-independent manner, indicating that NEDD8 binding to the yeast proteasome allows substrate delivery and degradation, as long as there is a flexible initiation region for engagement.

**Figure S2:**
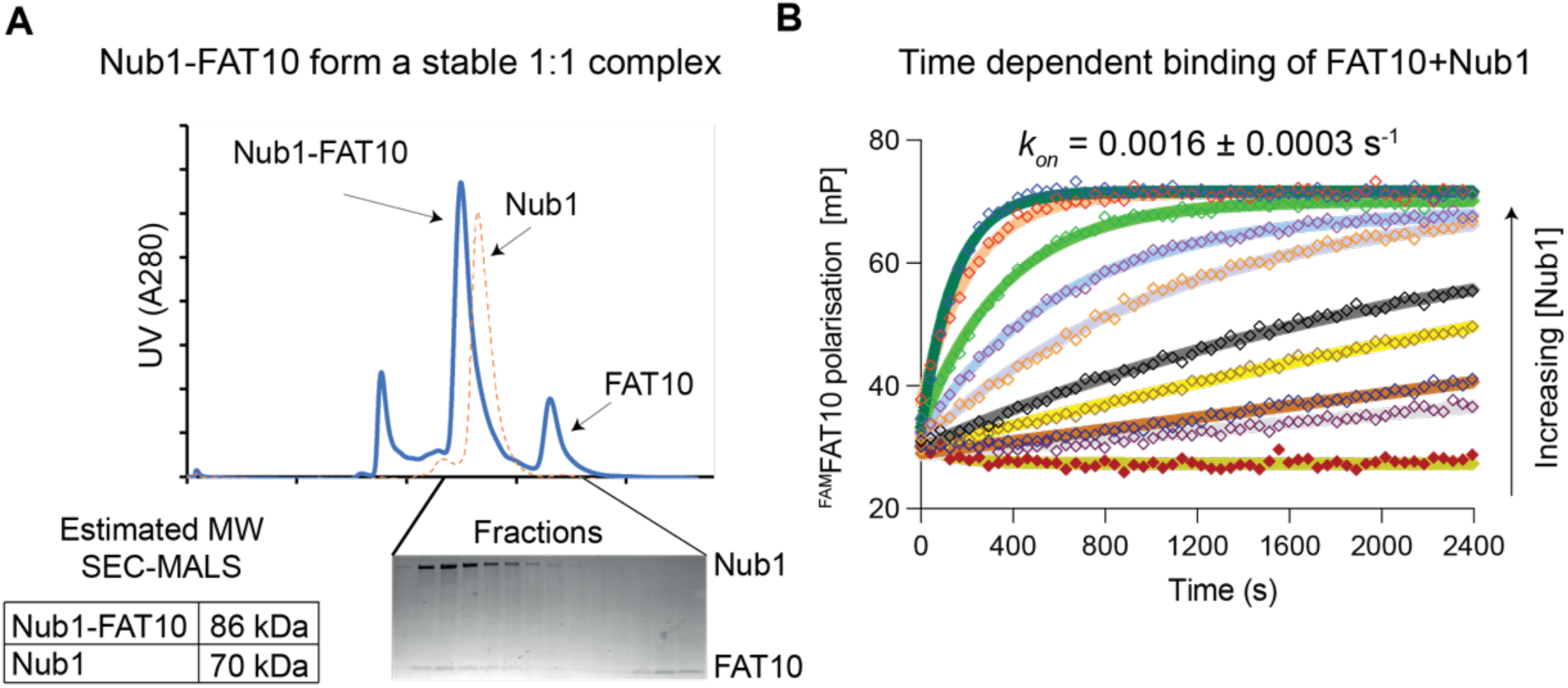
Nub1 and FAT10 slowly form a high-affinity 1:1 complex. **A**) Elution profiles for the size-exclusion chromatography of Nub1 alone (dotted orange line) or a sample in which Nub1 (50 μM) was preincubated with FAT10 (75 μM) for 60 min. The SDS-PAGE gel below shows samples of the individual fractions. The void peak likely originates from unfolded FAT10 and subsequent aggregation. **B)** Time courses for the fluorescence polarization of ^FAM^FAT10 (20 nM) after mixing with Nub1 at increasing concentrations (0, 16, 32, 63, 125, 250, 500, 1000, 2000 and 4000 nM).

**Figure S3:**
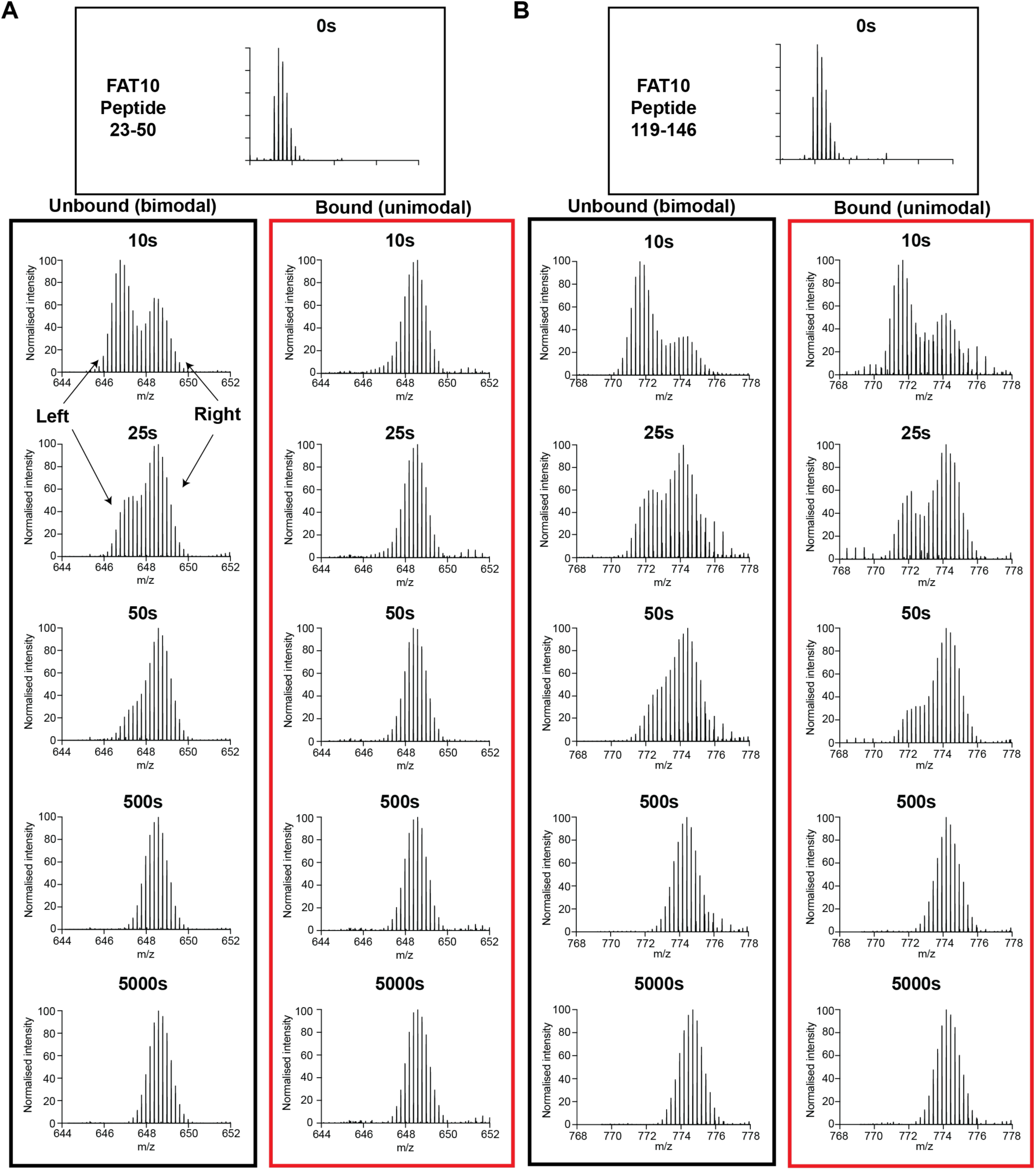
Example mass spectra from HDX-MS experiments of FAT10 versus FAT10 in the presence of excess Nub1. Intensity-normalized mass spectra for example peptides in FAT10’s UBL1 (**A**) and UBL2 domain (**B**) show bimodal deuterium uptake with two peaks, a left and a right distribution. The right distribution is fully exchanged, whereas the left peak shifts to the right and, in addition, its ratio relative to the right peak lowers over time. This behavior indicates that the peptide exchanges with a combination of EX1 kinetics (leading to a change in the peak ratio) and EX2 kinetics (leading to an average peak movement from left to right). The bimodal nature suggests that there are multiple species of FAT10, folded and unfolded, which transition on the order of minutes based on the EX1 kinetics of FAT10. In the presence of excess Nub1, the UBL1 peptide (A) shows unimodal behavior with the peptide signal shifted to the right, indicating a fully unfolded and fast exchanging species. In contrast, the UBL2 peptide (B) remains largely unaffected in its bimodal behavior by Nub1.

**Figure S4:**
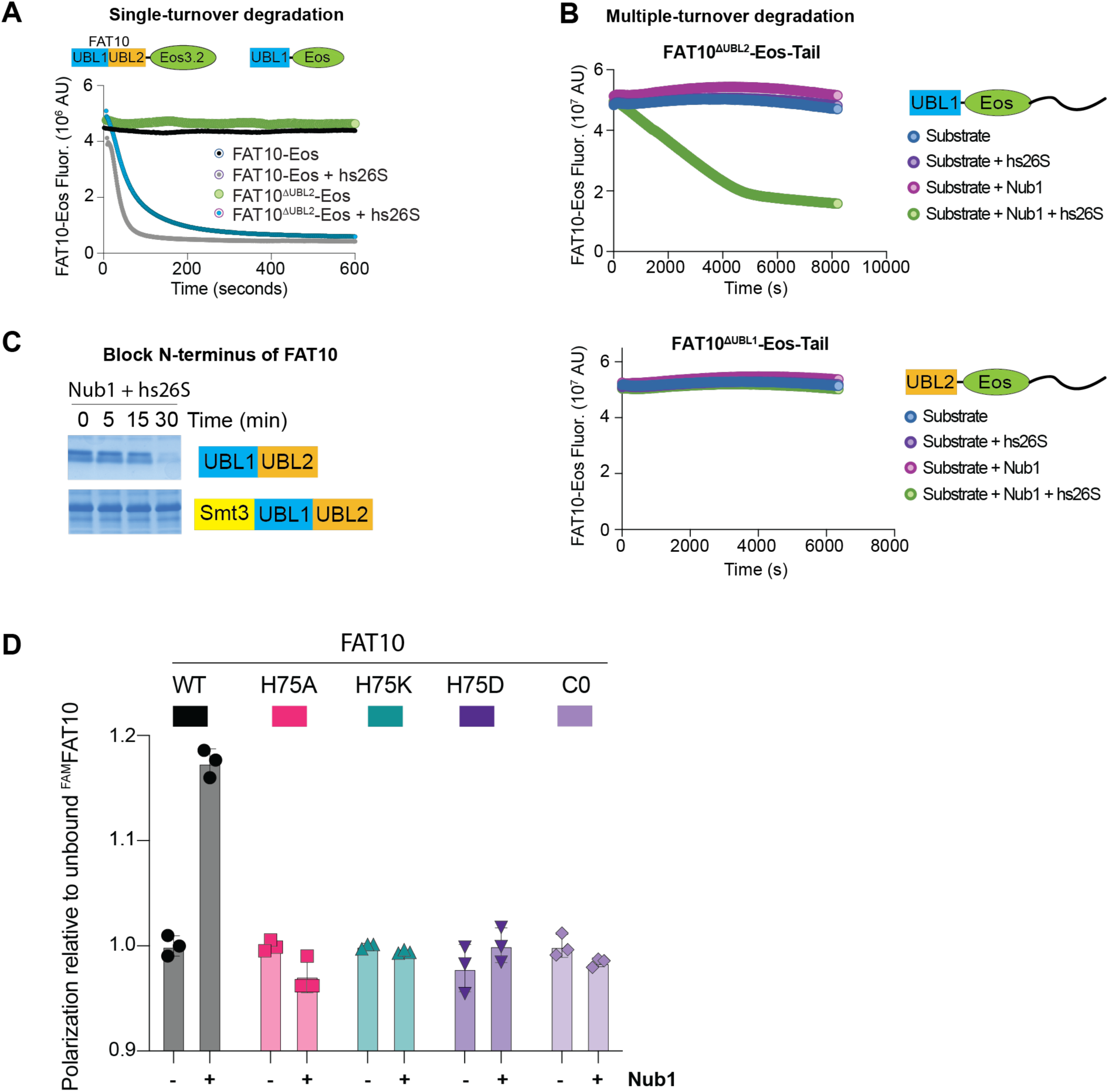
UBL1 is necessary and sufficient for Nub1-mediated degradation of FAT10. **A**) Single-turnover degradations of FAT10-Eos and FAT10^ΔUBL2^-Eos (100 nM) by *hs*26S proteasome (2 μM) in the presence of Nub1 (10 μM) indicate that FAT10’s UBL1 is sufficient for degradation, but the UBL2 domain is likely involved in forming a more productive complex with the proteasome for FAT10 turnover. **B)** Fluorescence-based assay for the degradation of FAT10^ΔUBL2^-Eos-tail (5 μM, top) and FAT10^ΔUBL1^-Eos-tail (5 μM, bottom) by *hs*26S proteasome (100 nM) in the absence and presence of Nub1 (15 μM). Nub1 and FAT10’s UBL1 domain are required for FAT10 delivery and degradation, even in the presence of a long initiation region. **C)** Coomassie-stained gel to analyze the degradation of FAT10 or a His-Smt3-FAT10 fusion (10 μM) by *hs*26S proteasome (100 nM) in the presence of Nub1 (5 μM). The lack of His-Smt3-FAT10 degradation indicates that FAT10’s free N-terminus is critical for insertion into and engagement by the proteasomal ATPase motor. **D)** Fluorescence polarization measurements analyzing the complex formation between wild-type or mutant ^FAM^FAT10 (100 nM) and Nub1 (10 μM), which were mixed and incubated for 30 mins on ice prior to the measurements. Shown are the mean values and standard deviations for N = 3 technical repeats, normalized to the values for free ^FAM^FAT10.

**Figure S5:**
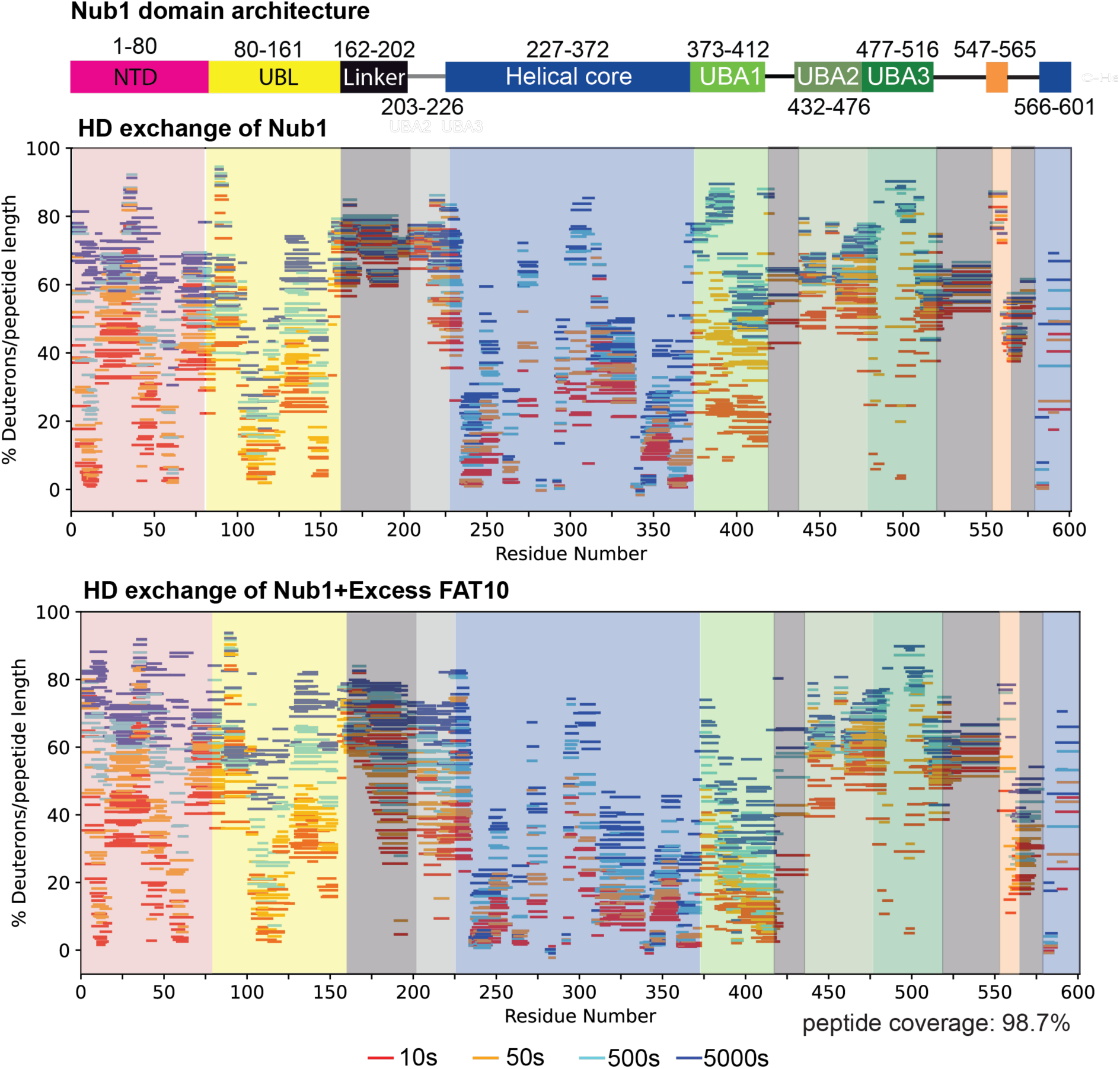
Woods plots for the deuteration levels of Nub1 in the absence or presence of excess FAT10. Displayed are the deuteration levels in percent for each peptide found in both Nub1 alone and Nub1 with excess FAT10 at time point of 10 – 5000 s after mixing with D_2_O. Each peptide is positioned in alignment with the amino acid sequence of Nub1 on the X-axis and the domain schematic above. The y-axis shows deuteration levels in percent relative to the theoretical minimum and maximum amide-proton exchange for each peptide.

**Figure S6:**
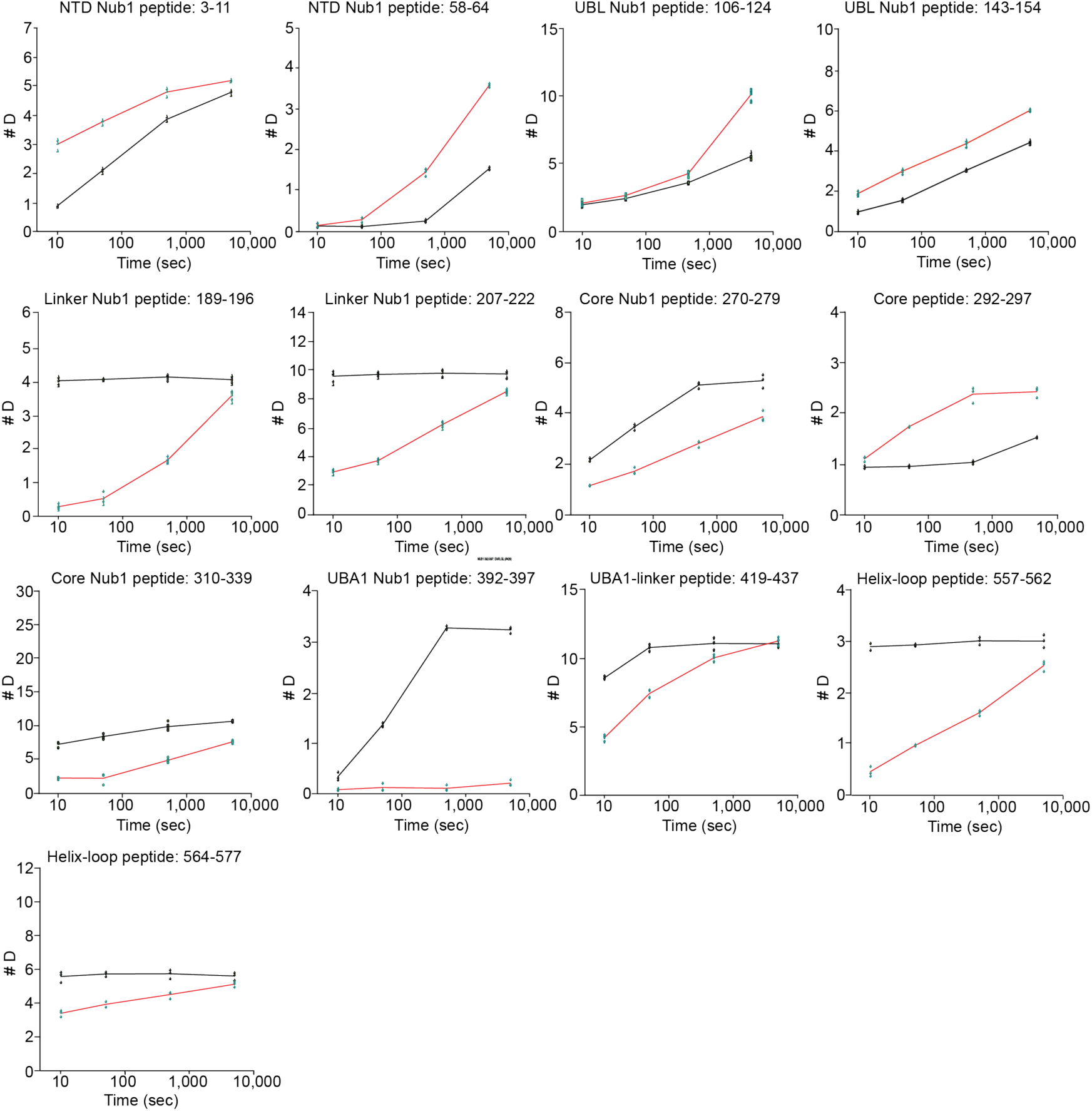
Deuterium uptake plots for selected Nub1 peptides. Plots show the deuterium uptake at 10, 50, 500 and 5000 s for individual peptides of Nub1 (black curves) compared to Nub1 in the presence of FAT10 (red curves).

**Figure S7:**
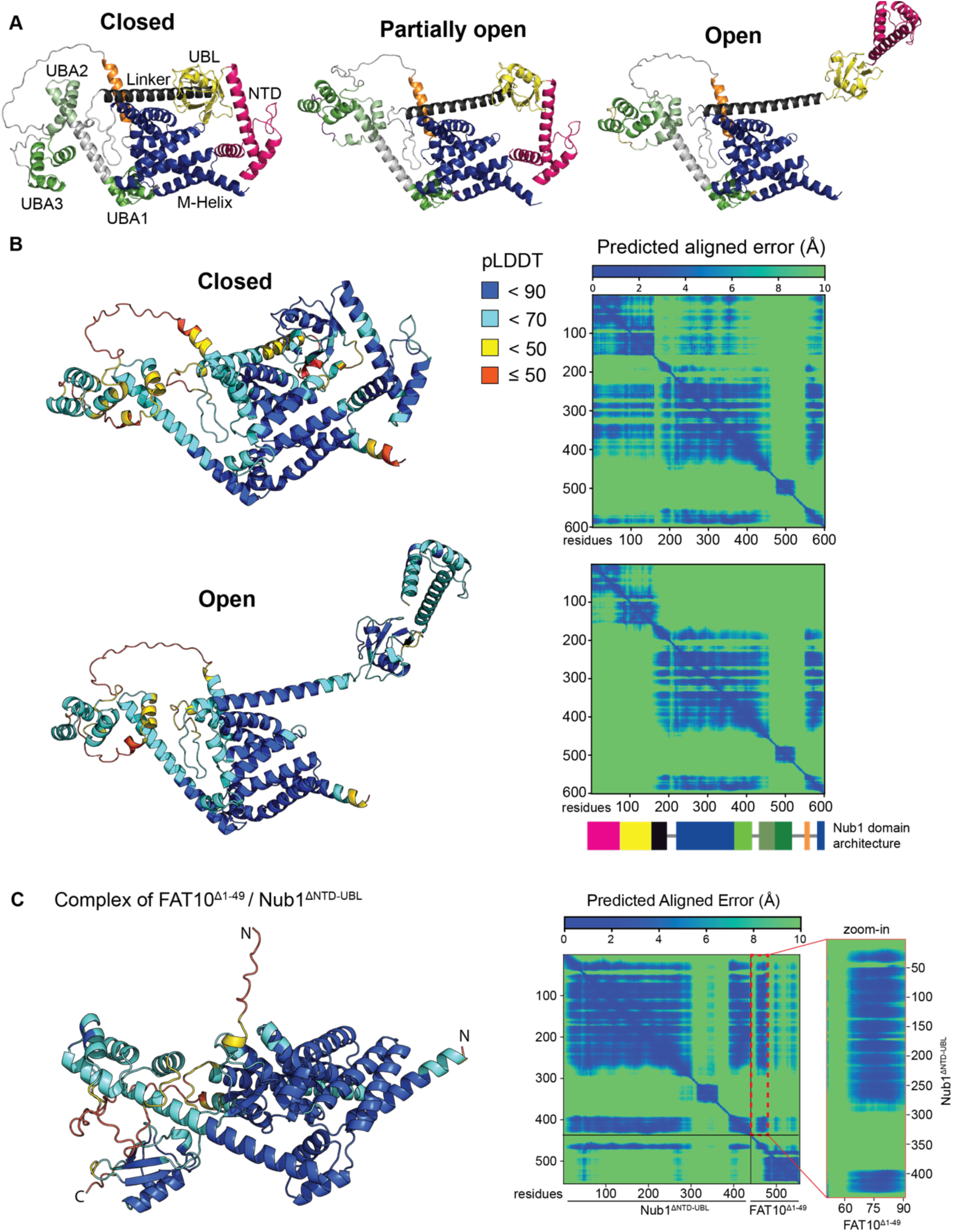
AlphaFold scores for selected Nub1 model predictions. **A**) The google notebook of Collabfold ^76^, which is based on the AlphaFold algorithm, was used to generate 5 models of Nub1. Models with notable differences were selected and named ‘open’, ‘partially open’, and ‘closed’ based on the position of Nub1’s ubiquitin-like (UBL) domain. Models are colored as in Figure 3B, with the N-terminal domain (NTD) in pink, the UBL domain in yellow, the helical linker in black, the disordered loop in grey (referred to as the beta-strand linker or BS-Trap), the core body in blue, and the three Ubiquitin-associated (UBA) domains in shades of green. **B)** AlphaFold models for the ‘closed’ state (top left) and ‘open’ state (bottom left) of Nub1 are colored based on predicted Local Distance Difference Test (pLDDT) scores. In brief, scores above 90 are considered as very accurate for local features and potentially side chains, 70-90 as good for overall fold with secondary structure, 50-70 as low confidence including the visual representation, and below 50 as likely disordered until in the right context. Right: Predicted aligned errors up to 10 Å for the closed and open states of Nub1, which describe the confidence in the relative positions of two residues and in domain-to-domain packing within the predicted structures. **C)** Left: AlphaFold-multimer model for the Nub1/FAT10 complex colored based on pLDDT scores. Right: predicted aligned error (PAE) depicts the quality of the AlphaFold-multimer predictions for domain docking and protein-protein interactions. The zoomed-in section for residues 50-92 of FAT10^Δ1–49^ highlights the confidence in the interaction between Nub1 and the last beta strand in FAT10’s UBL1 domain.

**Figure S8:**
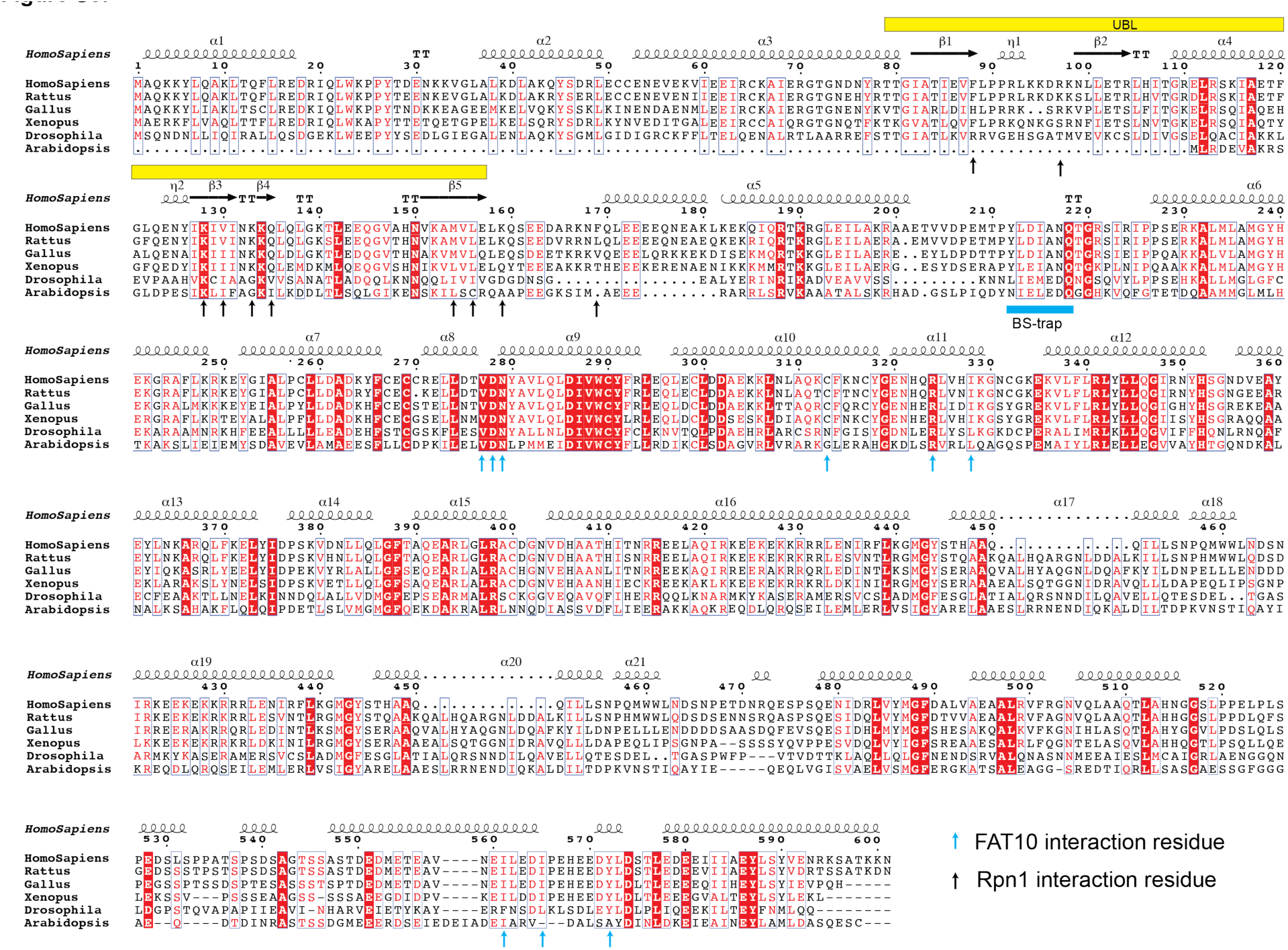
Multiple sequence alignment of Nub1 amino acid sequences from model organisms. Nub1 is conserved in evolution from mammals to plants, unlike FAT10 which is only found in mammals. Clustal omega was used to align the Uniprot sequences, and the schematic was generated with ESPript ^77^ (https://espript.ibcp.fr) based on the AlphaFold model of Nub1 and input multiple sequence alignments.

**Figure S9:**
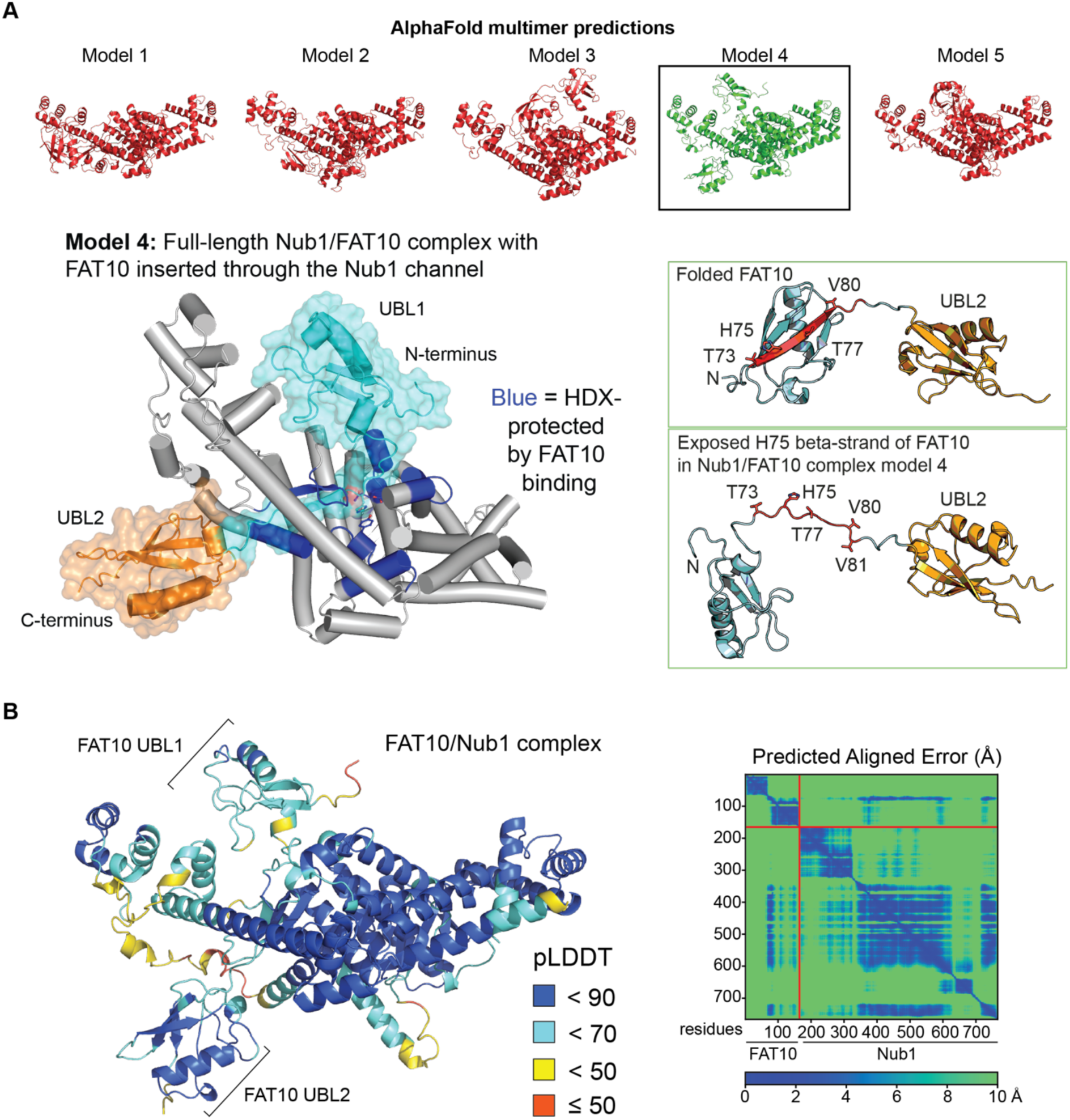
AlphaFold-multimer model of the full-length Nub1/FAT10 complex. **A**) Top: AlphaFold-multimer generated a range of models with FAT10 in various positions. Most were not confident based on the predicted aligned error (PAE) or not consistent with our results from biochemical and HDX-MS experiments. However, one model was consistent with those data and showed FAT10 inserted through the Nub1 channel, with the H75 beta-strand of FAT10 forming an anti-parallel beta sheet with beta-strand (BS)-trap linker of Nub1 (bottom left), similar to the structural model obtained with truncated Nub1 and FAT10 (Figure 3C). Bottom right: Comparison of folded FAT10 and FAT10 in the Nub1/FAT10 structural model 4 with unfolded UBL1 domain and exposed H75 beta strand. **B)** Left: pLDDT scores mapped onto the full-length Nub1/FAT10 complex structural model 4. Right: PAE scores for the complex. Of note, the UBL domain of Nub1 remains docked to the core body of Nub1 in this model, however, this is inconsistent with our HDX-MS data that suggest release and exposure of this UBL domain upon FAT10 binding. This is likely an artifact of AlphaFold-multimer, which adds weights to domain-domain and protein-protein interactions, giving rise to predictions where interfaces might be correct, but not necessarily in the right context. Nub1’s UBL domain likely binds to and releases from the core domain, and this equilibrium is shifted in the presence of FAT10. In addition, a partial beta-grasp fold is predicted for FAT10’s UBL1 domain, which may exist at times, but HDX-MS suggest a completely labile protein. The UBL1 domain of Nub1-bound FAT10 most likely lacks stable secondary structure elements that would normally slowly exchange in HDX experiments.

**Figure S10:**
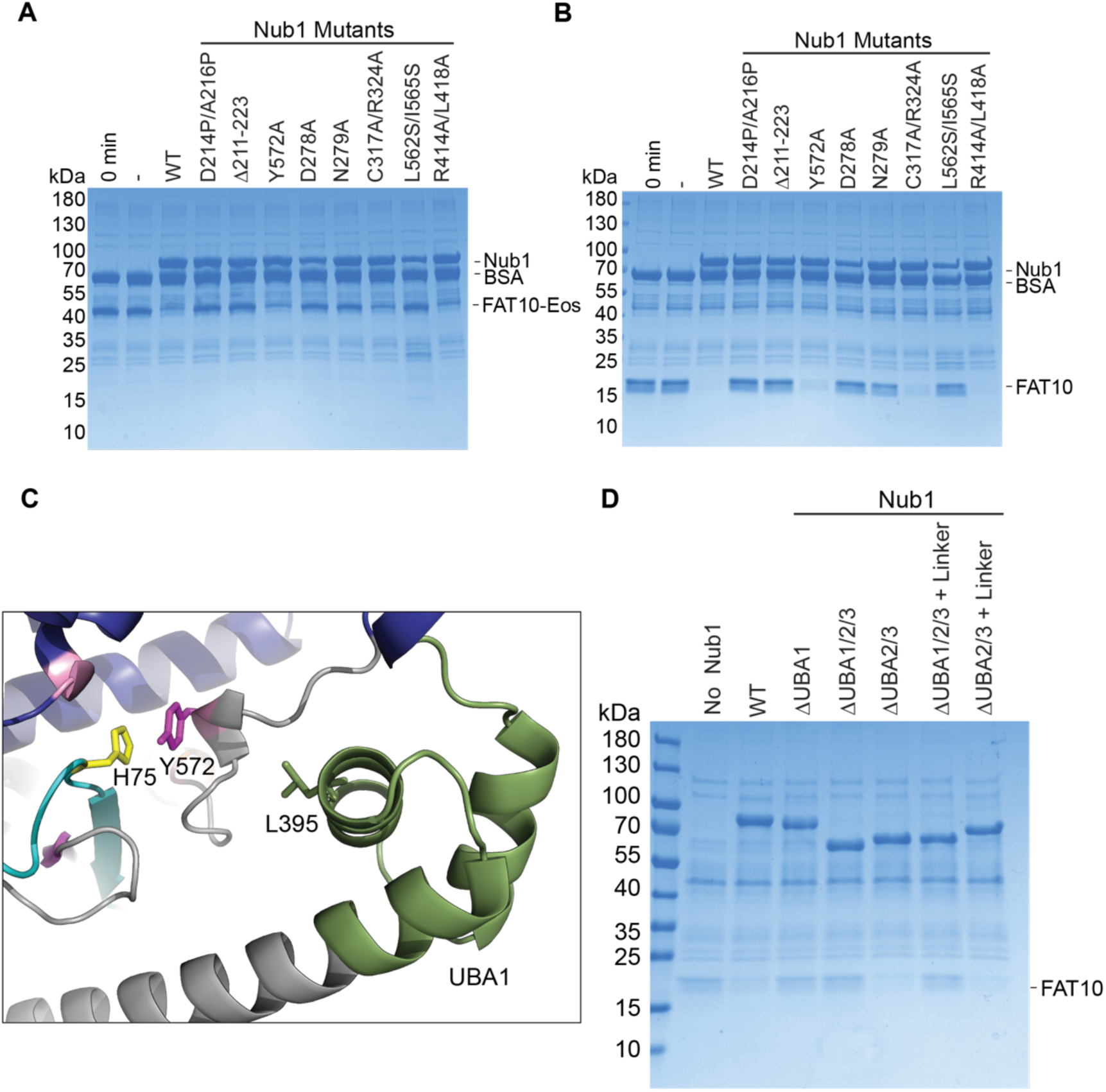
NUB1’s UBA1 domain and BS-Linker are critical for FAT10 degradation by the *hs*26S proteasome. **A**) Coomassie-stained SDS-PAGE gel analysis of the endpoints for the degradation of FAT10-Eos (1 μM) by the *hs*26S proteasome (100 nM) in the presence of various Nub1 mutants (5 μM). **B)** Assay as in A), but using FAT10 (10 μM). **C)** Structural model of the Nub1/FAT10 complex, with FAT10 in cyan and Nub1 in blue, grey, and green, showing the direct interaction of FAT10’s H75 with Nub1’s Y572, which appears to be stabilized by L395 and a helix in NUB1’s UBA1 domain. This interaction explains the FAT10-binding-induced protection of Nub1’s UBA1 domain in HDX-MS experiments. **D)** Coomassie-stained SDS-PAGE analysis of FAT10 degradation (5 μM) by *hs*26S proteasome (100 nM) in the presence of various truncation mutants of Nub1 (10 μM), showing that Nub1’s UBA2 and UBA3 domains are dispensable, while the UBA1 domain is essential for degradation.

**Figure S11:**
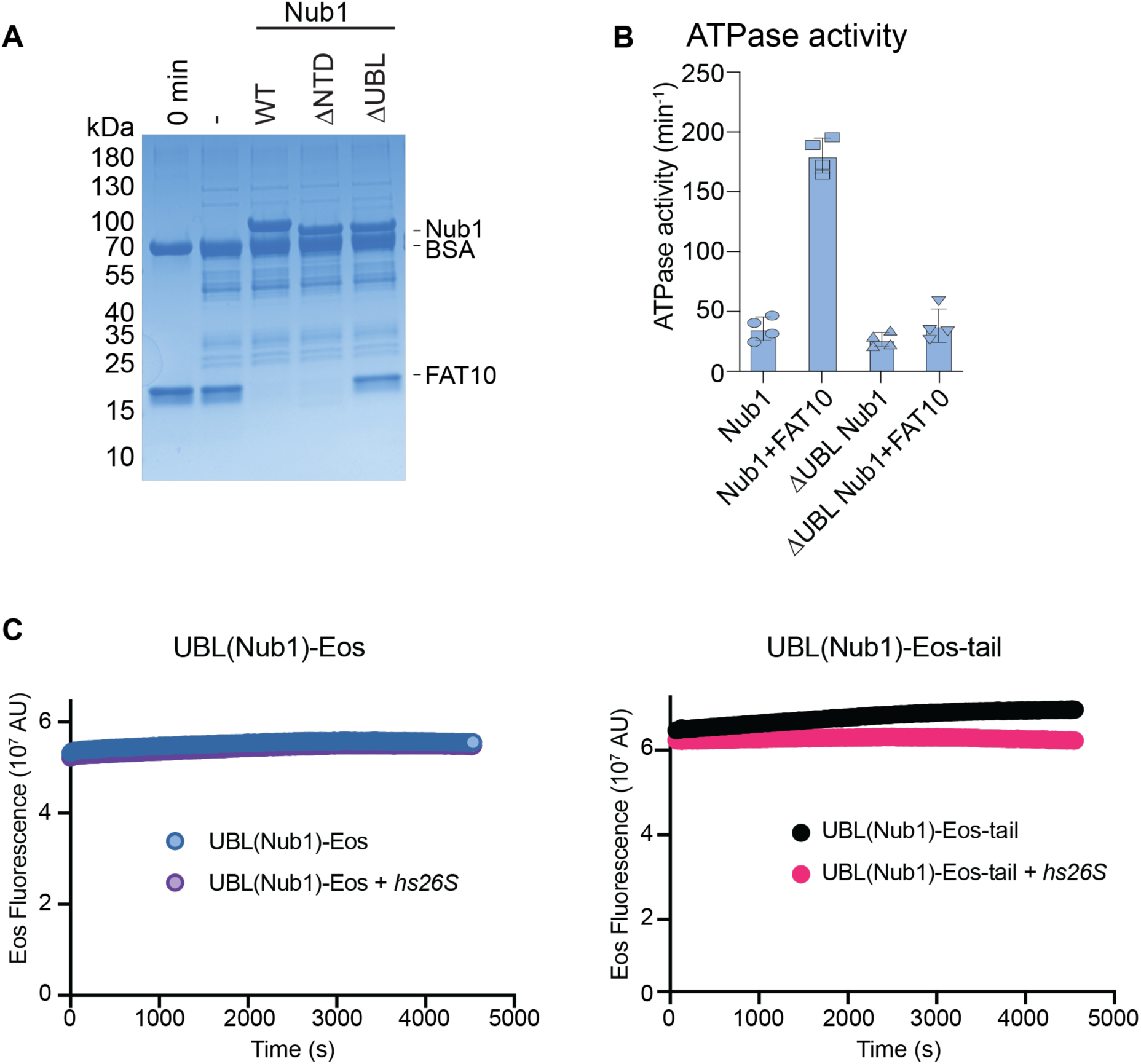
Nub1’s UBL domain is critical for FAT10 delivering to the proteasome. **A**) Coomassie-stained SDS-PAGE gel analysis of the endpoints for the degradation of FAT10 (10 μM) by *hs*26S proteasome (100 nM) in the presence of various Nub1 truncation mutants (2.5 μM). **B)** ATPase activity of *hs*26S proteasome in the absence and presence of Fat10, Nub1, or Nub1^ΔUBL^. Stimulation of ATP hydrolysis depends on Nub1’s UBL domain for FAT10 delivery and the engagement of the FAT10 with the ATPase motor to shift the proteasome conformation from engagement-competent to processing states. **C)** Nub1’s UBL domain is not sufficient for delivering a substrate for degradation. Example fluorescence traces for the incubation of the UBL(Nub1)-Eos fusion substrate (5 μM, left) or the UBL(Nub1)-Eos-tail fusion substrate (5 μM, right) with *hs*26S proteasome (100 nM).

**Figure S12:**
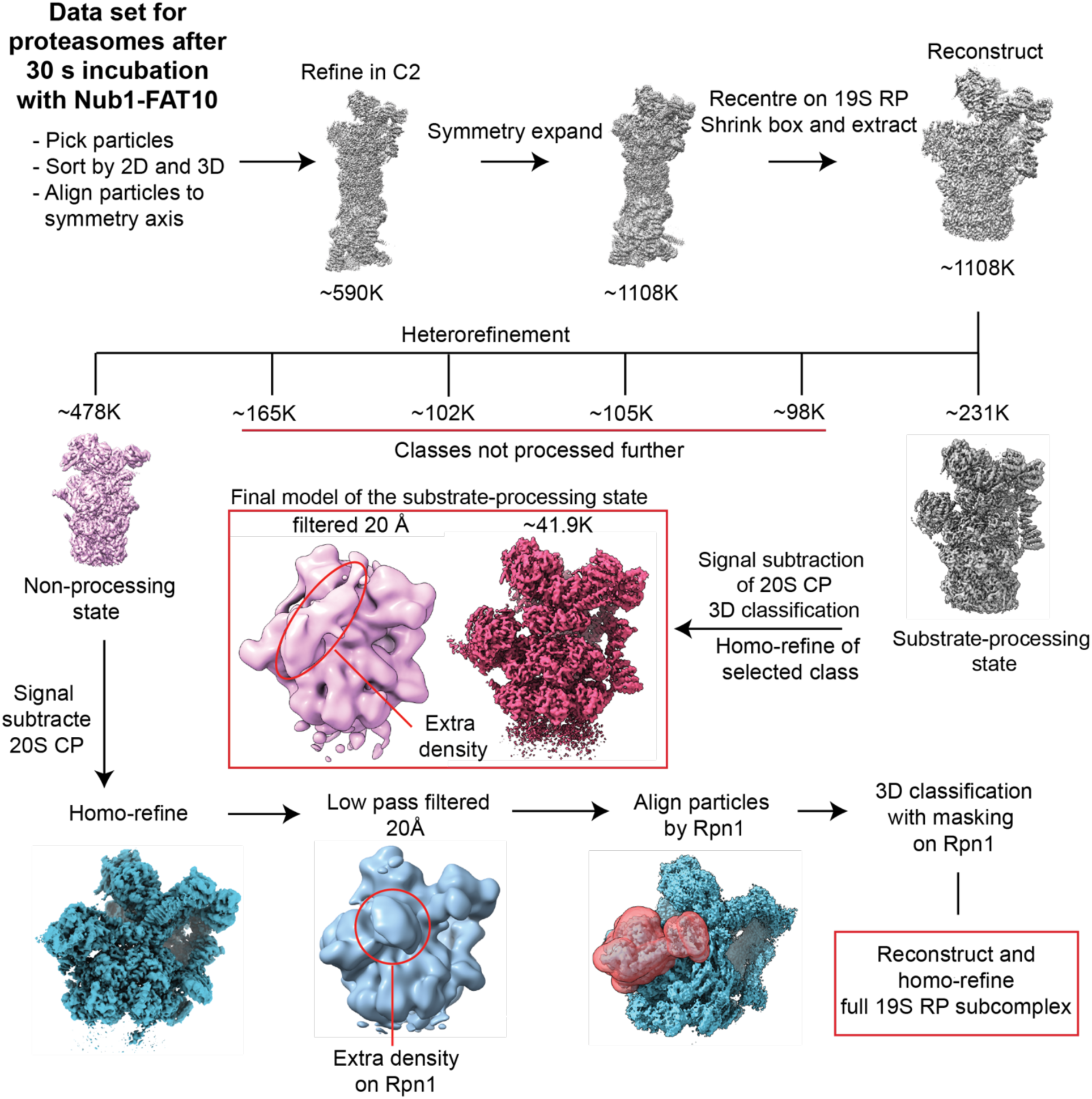
Processing workflow for the cryo-EM data set of proteasomes incubated with Nub1/FAT10 for 30 s. Preincubated FAT10-Eos (5 µM) and Nub1 (6 µM) were added to *hs*26S proteasome (2 µM) for 30 seconds before plunge freezing for Cryo-EM. The further classification of models for the non-processing proteasome is described in Figures S13 and S14. CryoSparc was used for all data processing and ChimeraX for visualization.

**Figure S13:**
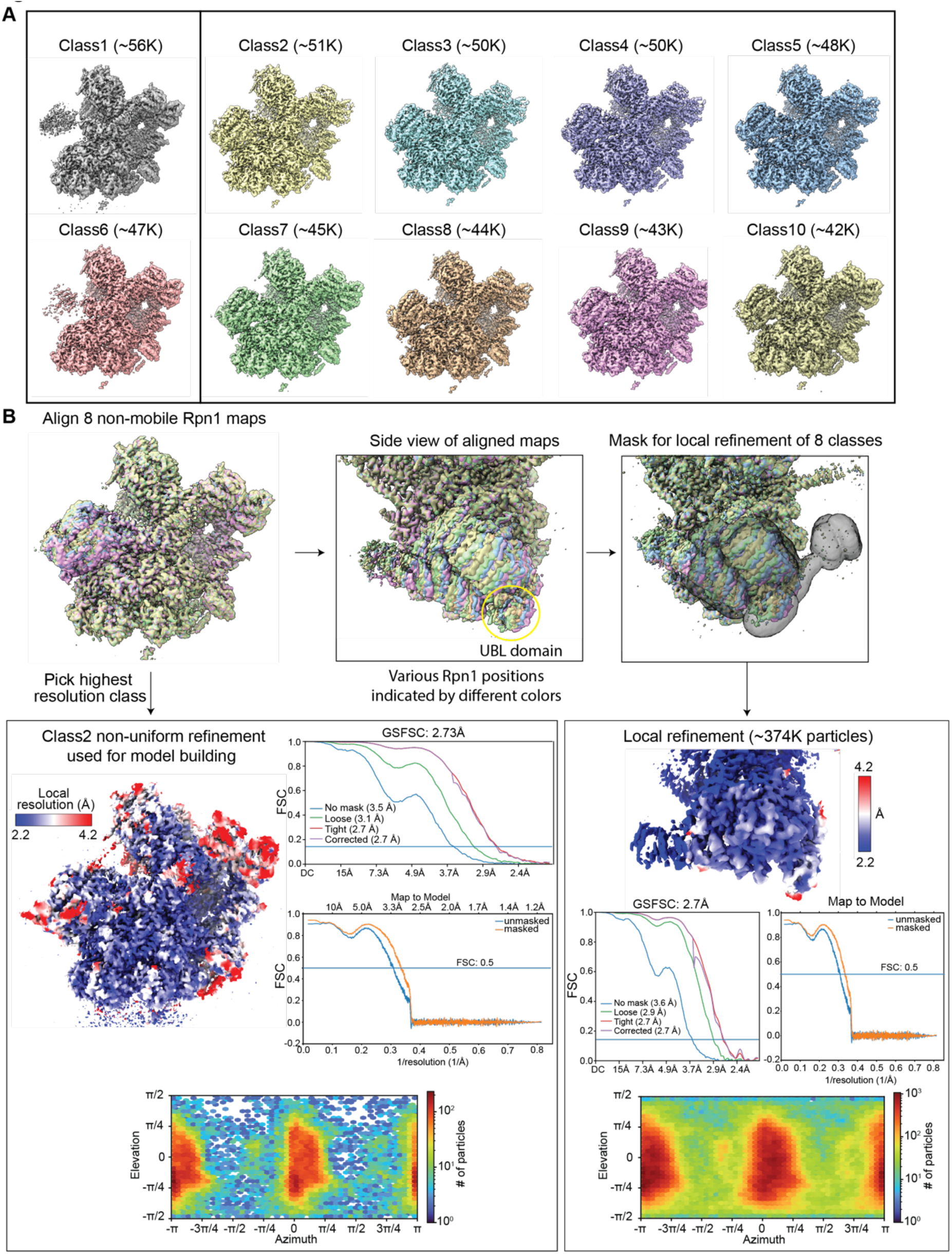
Continued cryo-EM data processing for the 30 s data set, focusing on Rpn1 classification. **A**) Using Rpn1-masked classification and refinement, 10 models of the non-processing *hs*26S proteasome are shown with particle numbers. Models 1 and 6 (left) show a blurry Rpn1 with continuous density, while the other 8 exhibit a resolved Rpn1 yet in slightly varied positions. **B)** An alignment of all Rpn1-resolved classes highlights these subtle differences in Rpn1 and therefore Nub1’s UBL position. The highest resolution class of the 19S RP subcomplex was further refined and used to build an atomic model (bottom left), for which the Gold-Standard FSC plot, map-to-model FSC plot, and distribution of particle orientations are show. Since Nub1’s UBL domain was at slightly lower resolution, further local refinement was performed on the Rpn1-Nub1 UBL interaction with all 8 Rpn1-resolved classes (bottom right), also showing the Gold-Standard FSC plot, map-to-model FSC plot, and distribution of particle orientations.

**Figure S14:**
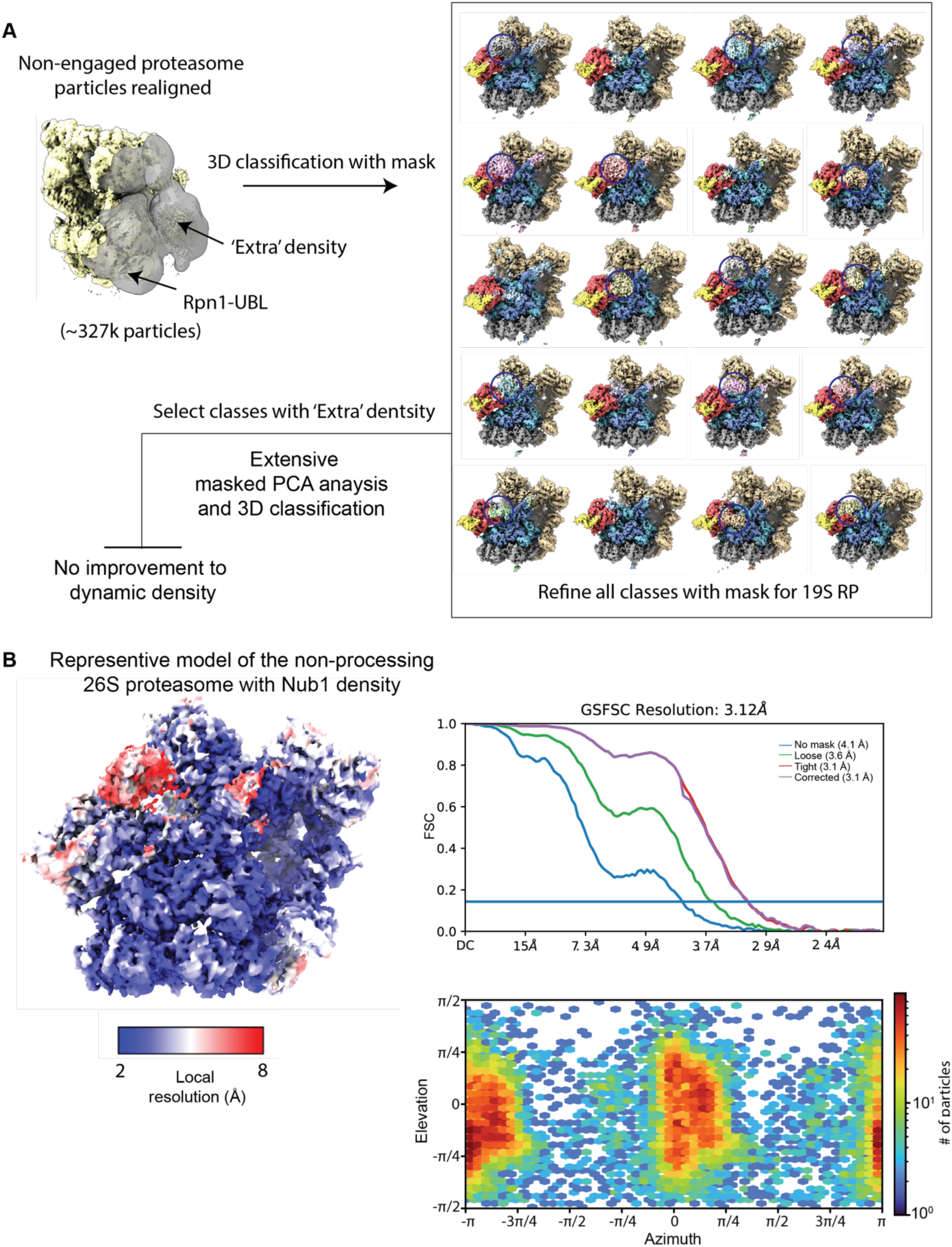
Continued Cryo-EM processing for the 30 s data set, focusing on the extra density connected to Nub1’s UBL domain. **A**) Extensive attempts were made to resolve the additional density attached to Nub1’s UBL domain. The ‘best’ results for representation were low-resolution amorphous masses for the extra density, which likely contains Nub1’s core domain bound to an unfolded FAT10-Eos molecule. This suggests that Nub1 is dynamic with respect to the relative orientation of its core and UBA domains and is moving continually relative to its UBL domain, independent of the 26S proteasome. Focused refinements before and after particle subtractions of the 19S RP signal for extra density also failed to generate any interpretable models, likely to due to weak signal compared to the large 26S proteasome, the continuous motions within Nub1 itself, and the unfolded FAT10 molecule which in intrinsically dynamic. **B)** Representative model for the non-processing proteasome with bound Nub1/FAT10-Eos. Shown also are the Gold-Standard FSC plot and distribution of particle orientations.

**Figure S15:**
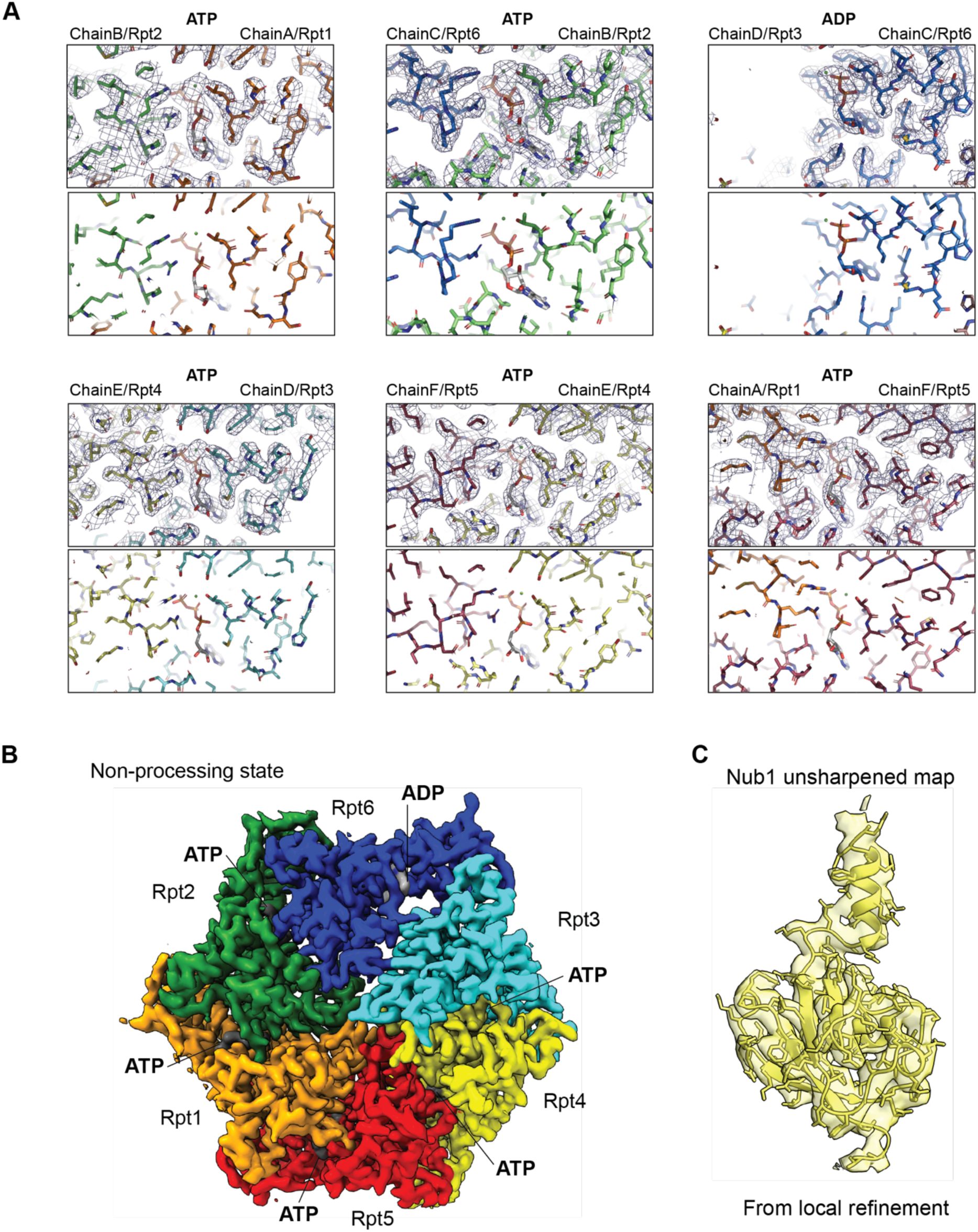
Examples of the well-resolved density for the ATPase motor and Nub1’s UBL domain from the 30 s data set of the non-processing *hs*26S proteasome. **A**) Atomic model focused on the nucleotide binding sites of Rpt1 – Rpt6 in unsharpened maps generated by non-uniform refinement of Class2 non-processing proteasomes. The name of each chain/ATPase subunit and its clockwise-next neighbor in the ATPase ring are shown above the maps on the right and left side, respectively. Rpt1 is depicted in orange, Rpt2 in green, Rpt6 in blue, Rpt3 in cyan, Rpt4 in yellow, and Rpt5 in red. **B)** Top view of ATPase motor density colored as in A) with nucleotide positions and identity indicated. **C)** Nub1’s UBL domain density derived from a locally refined model.

**Figure S16:**
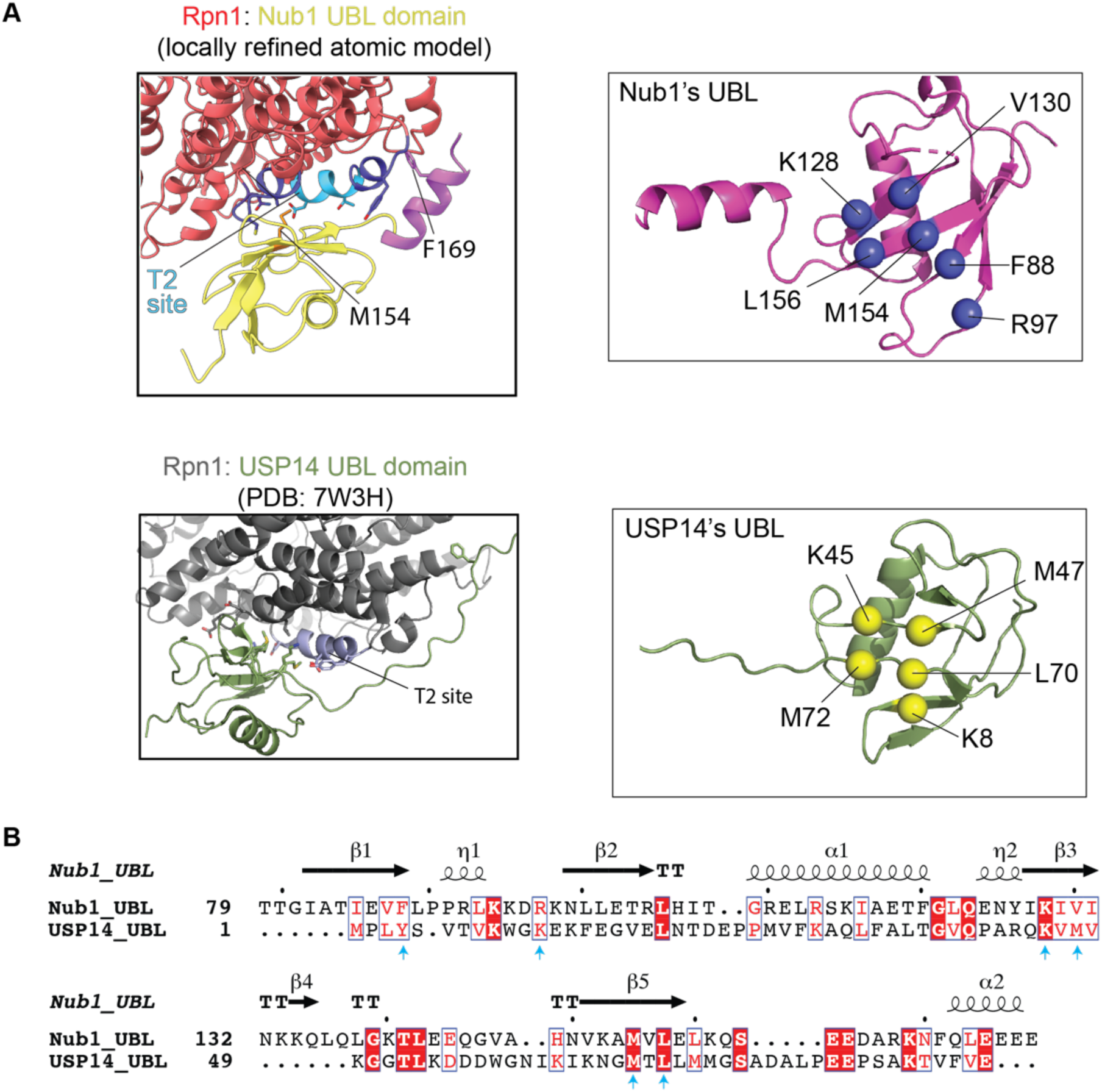
Comparison of Nub1’s UBL domain and USP14’s UBL domain bound to the T2 site of Rpn1. **A**) Left top and bottom: Structures of Nub1’s UBL domain (yellow) and USP14’s UBL domain (green) bound to Rpn1’s T2 site (light and slate blue). The representations highlight how both UBL domains bind the T2 site of Rpn1 through their beta sheet surface, which is structurally equivalent to the ubiquitin I44 hydrophobic patch as a common site for protein-protein interactions. Both UBL domains also use part of a C-terminally located linker, which connects the UBL domain to the rest of the respective protein, for additional interactions with Rpn1, yet at distinct sites. The positions of Nub1’s UBL domain and USP14’s UBL domain slightly vary with respect to Rpn1, potentially due differences in proteasome conformations and transient interactions formed between Rpn1 and the linkers. **B)** Sequence alignment of the UBL domains of human Nub1 and human USP14, generated with Clustal Omega and ESPript ^77^ (https://espript.ibcp.fr).

**Figure S17:**
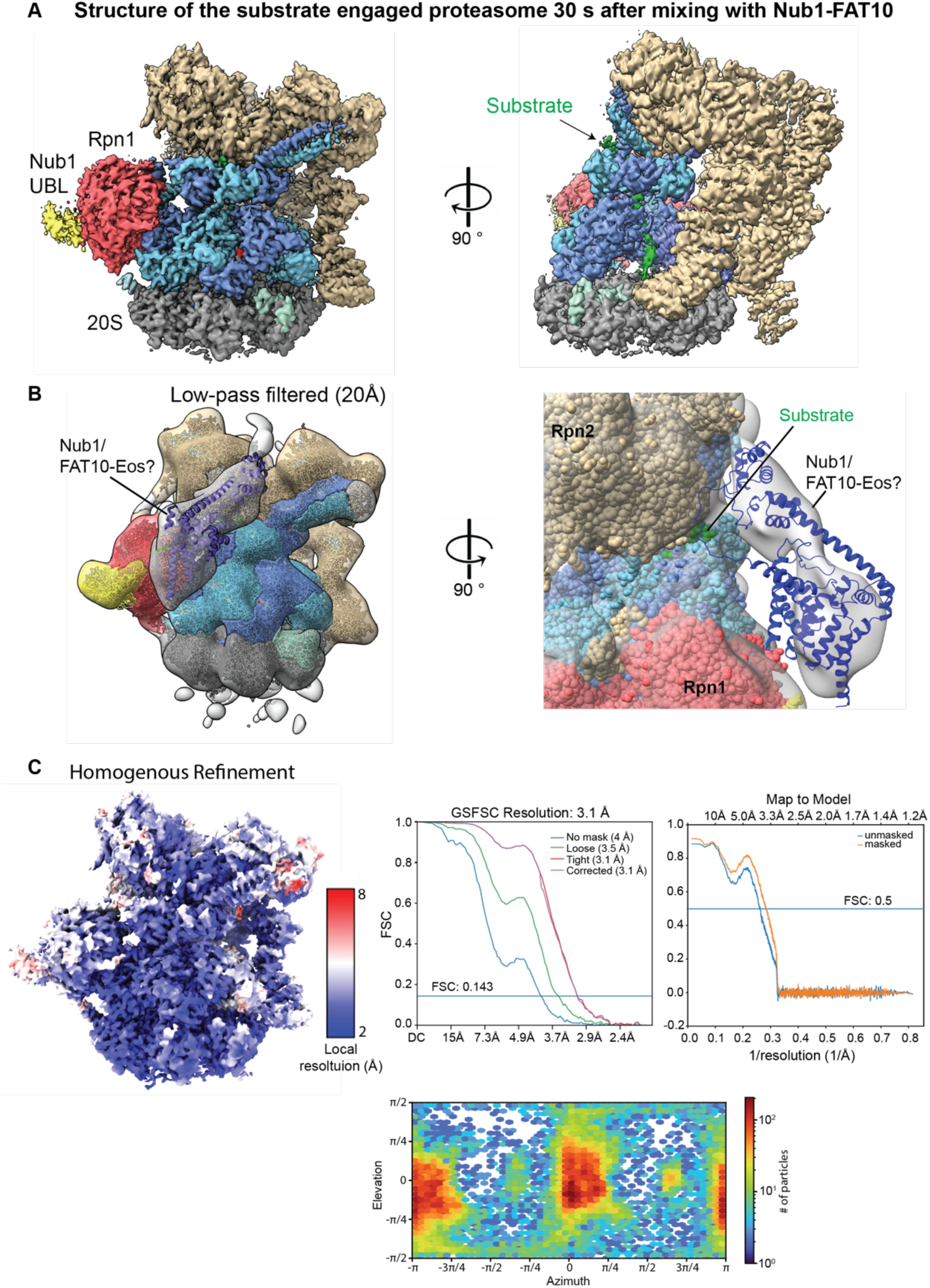
High-resolution model of the substrate-processing *hs*26S proteasome from the 30 s data set. **A**) Left: Reconstruction used for generating the atomic model, with Rpn1 shown in red, Nub1’s UBL domain in yellow, the ATPase motor in alternating blue and cyan, the 20S CP in gray, the lid subcomplex in sand color, and the substrate in green. Right: Reconstruction rotated by 90 degrees and with the density for the Rpt4 and Rpt5 ATPase subunits removed to visualize the translocating substrate density inside the central channel. **B)** A low pass filtered model of the density in A), shown for the entire top part of the proteasome (left) and focused on the space between Rpn1 and Rpn2 after a 90 degree rotation (right) to visualize the large extra density that connects to the well-resolved UBL domain of Nub1 and the translocating substrate. The size of this density agrees well with the size of Nub1, as indicated by the roughly docked structural model for Nub1’s core and UBA domains (purple ribbon presentation). However, Nub1 could not be well resolved, as it is likely present in multiple different positions and with variable orientations of its domains relative to each other. **C)** Local resolution, GSFSC curve, map-to-model FSC, and distribution of particle orientations for the substrate-processing model of the *hs*26S proteasome.

**Figure S18:**
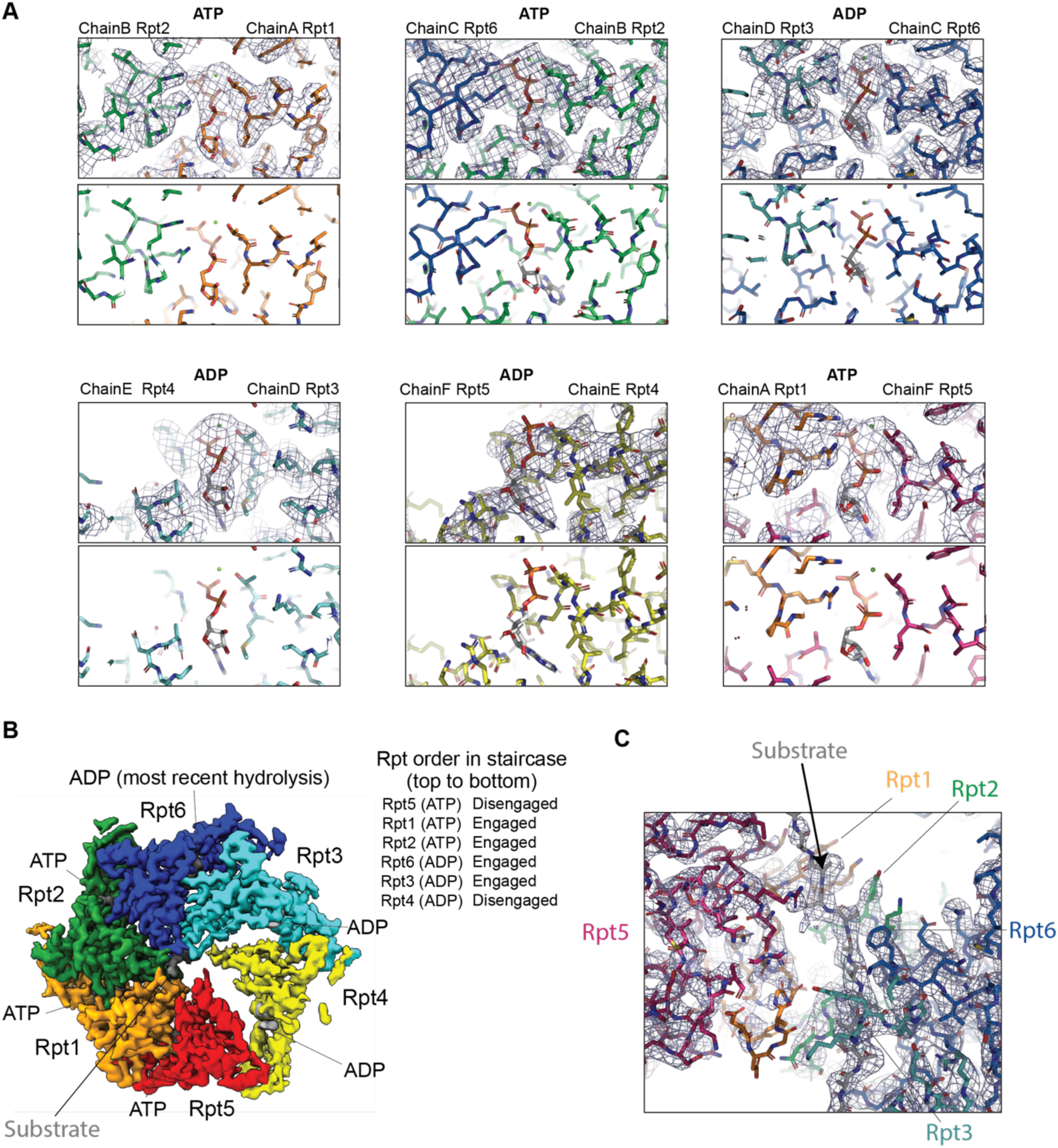
Examples of the well-resolved density for the ATPase motor and the translocating substrate from the 30 s data set of the processing *hs*26S proteasome. **A**) Atomic model focused on the nucleotide binding sites of Rpt1 – Rpt6 in unsharpened maps. The name of each chain/ATPase subunit and its clockwise-next neighbor in the hexameric ATPase ring are shown above the maps on the right and left side, respectively. Rpt1 is depicted in orange, Rpt2 in green, Rpt6 in blue, Rpt3 in cyan, Rpt4 in yellow, and Rpt5 in red. **B)** Top view of ATPase motor density colored as in A) with nucleotide positions and identity indicated. The Rpts are arranged in a spiral staircase, with Rpt5 at the top, Rpt3 at the bottom, and Rpt4 as the mobile seam subunits on its way to the top. Rpt1, 2, 6, and 3 are engaged with the substrate polypeptide, whereas Rpt4 is disengaged and Rpt5 has not yet engaged the substrate. The ADP in Rpt6 likely originated from the most recent ATP-hydrolysis event, which precedes the disengagement of Rpt3 from the substrate and the substrate re-engagement of Rpt4 at the top of the staircase. **C)** Side view of the translocating substrate polypeptide contacted within the central channel by the pore-1 loop tyrosine of multiple Rpt subunits.

**Figure S19:**
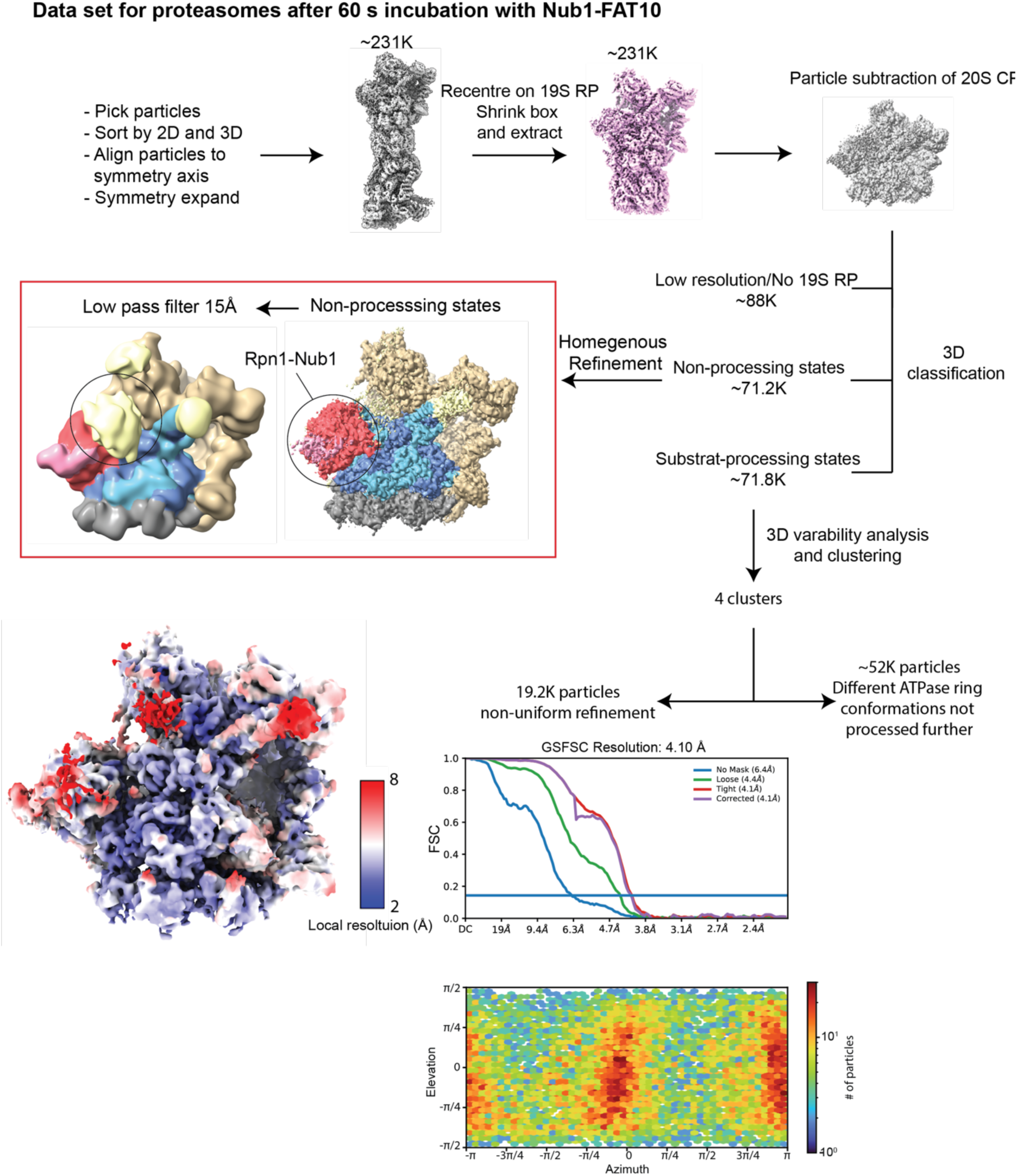
Cryo-EM data processing workflow for the 60 s data set of substrate-processing *hs*26S proteasome. A preincubated mixture of FAT10-Eos (5 µM) and Nub1 (6 µM) was added to *hs*26S proteasome (2 µM) for 60 s before plunge freezing for cryo-EM. The derived model shows an overall similar conformation as the substrate-processing state after 30 s incubation, albeit at lower resolution and with more density at the entrance to the central processing channel. Shown at the bottom are the local resolution, GSFSC curve, and distribution of particle orientations. CryoSparc was used for all data processing, with ChimeraX for visualization.

**Table S1:** Peptide-coverage statistics for HDX-MS experiments of FAT10 and Nub1. Peptides for FAT10 and Nub1 were only kept if found in both unbound and bound samples. Data were generated using HDExaminer 3.

|  | Peptides | Coverage | Average length | SD | Average Redundancy |
| --- | --- | --- | --- | --- | --- |
| Nub1 | 573 | 98.7% | 13.1 | 6.4 | 12.4 |
| FAT10 | 189 | 90.3% | 16.0 | 7.3 | 18.3 |

**Table S2:**
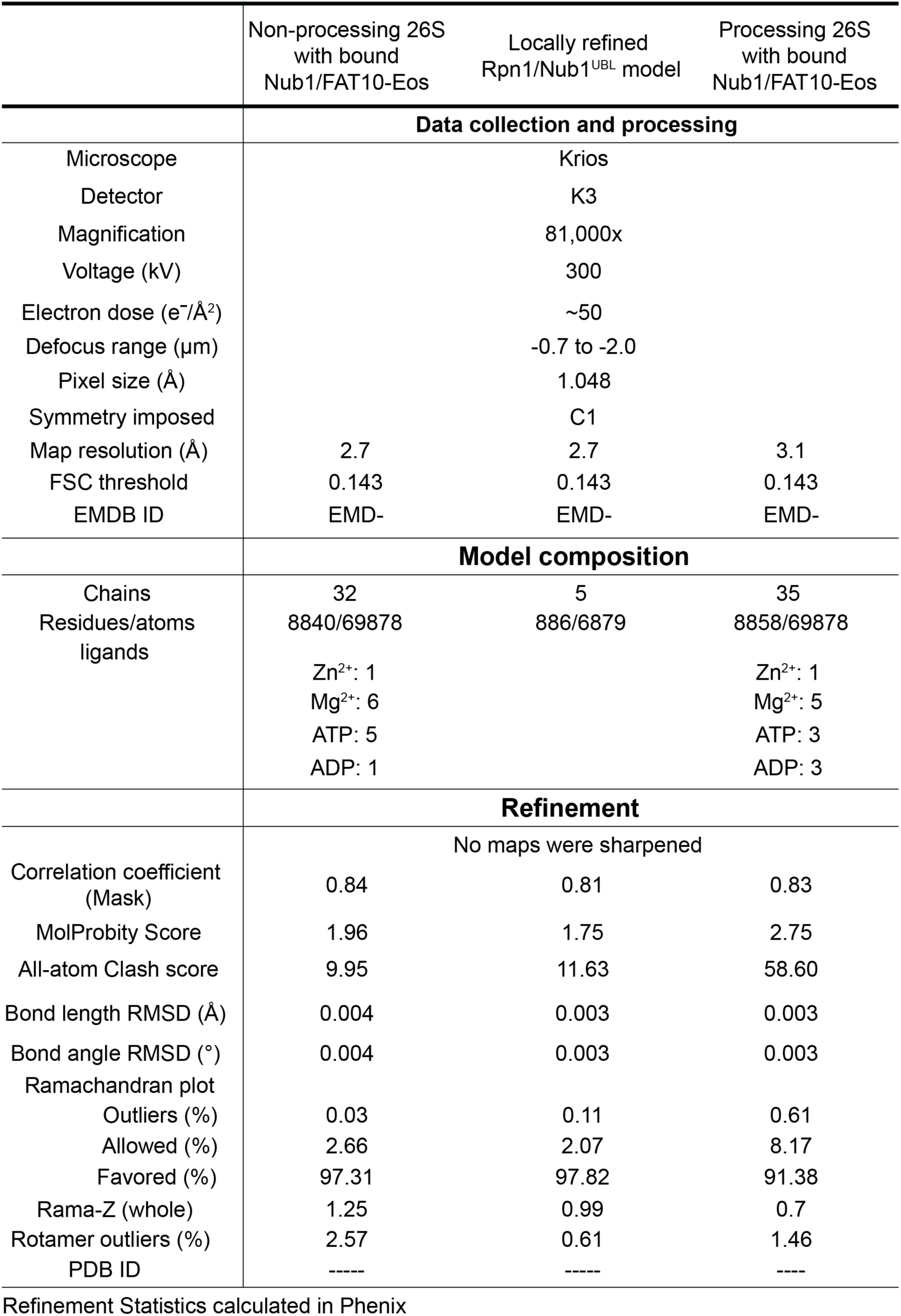
Cryo-EM data collection and model refinement statistics.

|  | Non-processing 26S<br>with bound<br>Nub1/FAT10-Eos | Locally refined<br>Rpn1/Nub1<sup>UBL</sup> model | Processing 26S<br>with bound<br>Nub1/FAT10-Eos |
| --- | --- | --- | --- |
| **Data collection and processing** |  |  |  |
| Microscope | Krios |  |  |
| Detector | K3 |  |  |
| Magnification | 81,000x |  |  |
| Voltage (kV) | 300 |  |  |
| Electron dose (e<sup>-</sup>/Å<sup>2</sup>) | ~50 |  |  |
| Defocus range (μm) | -0.7 to -2.0 |  |  |
| Pixel size (Å) | 1.048 |  |  |
| Symmetry imposed | C1 |  |  |
| Map resolution (Å) | 2.7 | 2.7 | 3.1 |
| FSC threshold | 0.143 | 0.143 | 0.143 |
| EMDB ID | EMD- | EMD- | EMD- |
| **Model composition** |  |  |  |
| Chains | 32 | 5 | 35 |
| Residues/atoms | 8840/69878 | 886/6879 | 8858/69878 |
| ligands | Zn<sup>2+</sup>: 1<br>Mg<sup>2+</sup>: 6<br>ATP: 5<br>ADP: 1 |  | Zn<sup>2+</sup>: 1<br>Mg<sup>2+</sup>: 5<br>ATP: 3<br>ADP: 3 |
| **Refinement** |  |  |  |
| No maps were sharpened |  |  |  |
| Correlation coefficient (Mask) | 0.84 | 0.81 | 0.83 |
| MolProbity Score | 1.96 | 1.75 | 2.75 |
| All-atom Clash score | 9.95 | 11.63 | 58.60 |
| Bond length RMSD (Å) | 0.004 | 0.003 | 0.003 |
| Bond angle RMSD (°) | 0.004 | 0.003 | 0.003 |
| Ramachandran plot |  |  |  |
| Outliers (%) | 0.03 | 0.11 | 0.61 |
| Allowed (%) | 2.66 | 2.07 | 8.17 |
| Favored (%) | 97.31 | 97.82 | 91.38 |
| Rama-Z (whole) | 1.25 | 0.99 | 0.7 |
| Rotamer outliers (%) | 2.57 | 0.61 | 1.46 |
| PDB ID | ---- | ---- | ---- |

Refinement Statistics calculated in Phenix

